# The *Staphylococcus aureus* small non-coding RNA IsrR regulates TCA cycle activity and virulence

**DOI:** 10.1101/2024.07.03.601953

**Authors:** Gustavo Rios-Delgado, Aubrey K. G. McReynolds, Emma A. Pagella, Javiera Norambuena, Paul Briaud, Vincent Zheng, Matthew J. Munneke, Jisun Kim, Hugo Racine, Ronan Carroll, Ehud Zelzion, Eric Skaar, Jeffrey L. Bose, Dane Parker, David Lalaouna, Jeffrey M. Boyd

## Abstract

*Staphylococcus aureus* has evolved mechanisms to cope with low iron (Fe) availability in host tissues. *S. aureus* uses the ferric uptake transcriptional regulator (Fur) to sense titers of cytosolic Fe. Upon Fe depletion, apo-Fur relieves transcriptional repression of genes utilized for Fe uptake. We demonstrate that an *S. aureus* Δ*fur* mutant has decreased expression of *acnA*, which codes for the Fe-dependent enzyme aconitase. Decreased *acnA* expression prevented the Δ*fur* mutant from growing with amino acids as sole carbon and energy sources. Suppressor analysis determined that a mutation in *isrR*, which produces a regulatory RNA, permitted growth by decreasing *isrR* transcription. The decreased AcnA activity of the Δ*fur* mutant was partially relieved by an Δ*isrR* mutation. Directed mutation of bases predicted to facilitate the interaction between the *acnA* transcript and IsrR, decreased the ability of IsrR to control *acnA* expression *in vivo* and IsrR bound to the *acnA* transcript *in vitro*. IsrR also bound to the transcripts coding the alternate TCA cycle proteins *sdhC*, *mqo*, *citZ*, and *citM*. Whole cell metal analyses suggest that IsrR promotes Fe uptake and increases intracellular Fe not ligated by macromolecules. Lastly, we determined that Fur and IsrR promote infection using murine skin and acute pneumonia models.

**Graphical Abstract:** 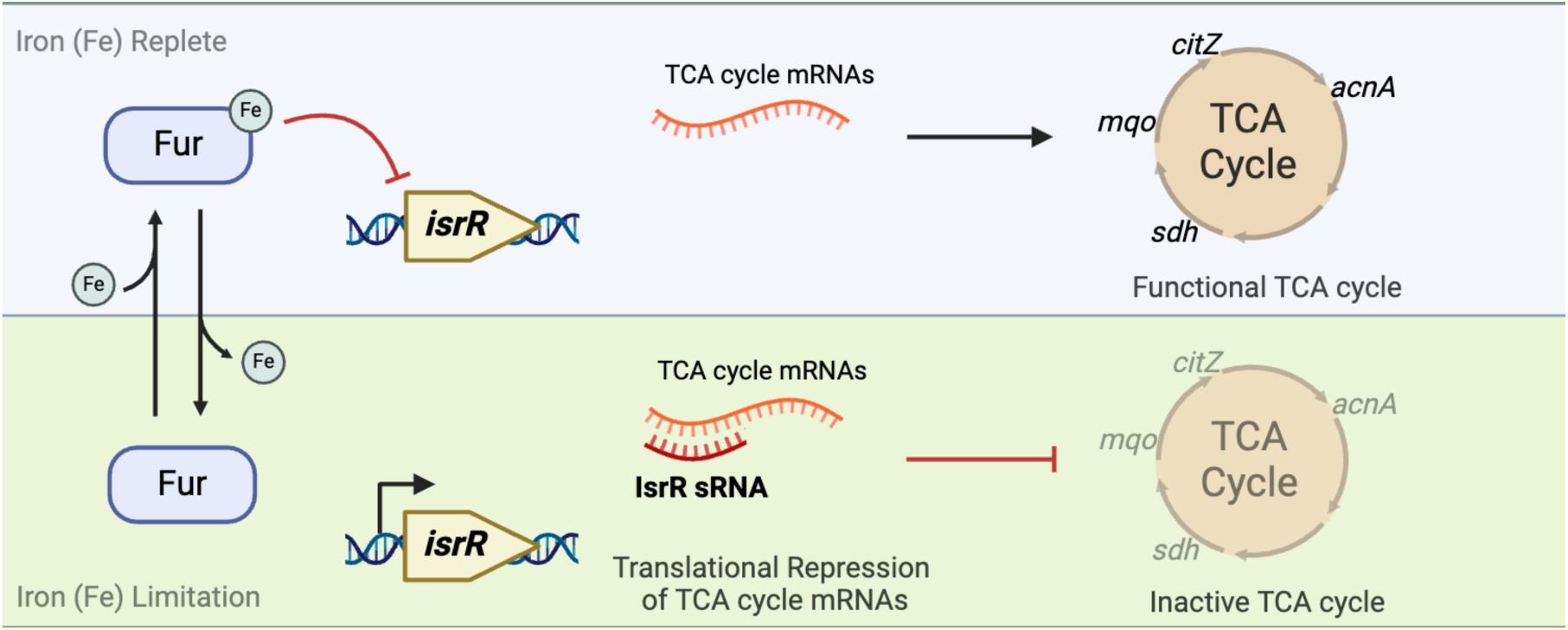

## Introduction

*Staphylococcus aureus* is an infectious agent and a primary cause of morbidity and mortality worldwide. It is commonly associated with community- and hospital-acquired infections, and it can cause a variety of ailments from minor soft tissue infections to more severe disease including septicemia and endocarditis (1). A subset of *S. aureus* associated infections are caused by strains resistant to commonly prescribed antibiotics complicating treatment (2).

The acquisition of ionic iron (Fe) is essential for *S. aureus*. *S. aureus* strains defective in the acquisition, uptake, or proper usage of intracellular Fe have decreased virulence in models of infection (3–6). The importance of Fe is highlighted by the fact that approximately 2% of the genomic protein coding open reading frames (ORFs) code for Fe acquisition systems. Iron functions as a cofactor to many proteins including iron-sulfur (Fe-S) cluster enzymes utilized for respiration and the tricarboxylic acid (TCA) cycle (7).

One strategy that mammals use to combat infections is to limit the availability of nutrients that are essential for bacterial growth. This process, known as nutritional immunity, limits bacterial access to trace metals including Fe ions (8). Inside the host, Fe is abundant, but it is typically found associated with proteins in prosthetic groups including heme or Fe-S clusters. Non-cofactor associated Fe in large part is ligated by proteins including transferrin, lactoferrin, and calprotectin (4,9,10). Therefore, within host tissues, the “free” or loosely ligated Fe concentrations are low as to restrict *S. aureus* growth (11). Hosts further limit Fe availability during infection by reducing the absorption of dietary Fe in a process referred to as anemia of inflammation (12). Pathogen containing macrophages export Fe ions and increase expression of ferritin to decrease free Fe titers (13,14). Individuals with the inborn disease hemochromatosis have increased Fe titers and an increased prevalence of infection highlighting the important of Fe limitation to prevent infection (15).

Given the scarcity of Fe in host tissues, bacteria must alter their gene expression profiles to colonize tissue and promote infection. A key part of the bacteria’s response to low Fe is the upregulation of Fe uptake systems including the expression of siderophores which are high affinity extracellular Fe ion chelating molecules that can compete with host proteins for Fe ions (16). Once bound to Fe, siderophores are transported back into the cell via specific surface receptors. Other strategies for Fe uptake include the use of cell-wall associated transferrin binding proteins, monovalent Fe uptake, and acquisition of Fe from host hemoglobin (17).

In *S. aureus*, the ferric uptake transcriptional regulator (Fur) senses and responds to cytosolic Fe titers (11). When associated with Fe, Fur typically acts as a transcriptional repressor (18,19). Growth in a low Fe medium promotes Fur demetallation, altered affinity for DNA, and expression of the Fur regulon. RegPrecise 3.0 predicts that there are 20 ORFs or operons in *S. aureus* that contain consensus Fur binding sites in their operators (20). In large part, these are genes involved in Fe uptake or storage. A *S. aureus* Δ*fur* mutant or growth during Fe limitation results in increased and decreased expression of glycolysis and the TCA cycle, respectively (18,21). The mechanism behind the Fur-dependent regulation of central metabolic pathways in *S. aureus* has remained elusive. In *Escherichia coli* and *Bacillus subtilis*, Fur directly controls the transcription of *ryhB* and *fsrA*, respectively (22,23). Both loci produce small non-coding regulatory RNAs (sRNA) that modulate gene expression by directly pairing with several messenger RNA (mRNA). In doing so, these sRNAs alter metabolism to spare Fe ions and redirect usage to prioritize essential processes. A recent publication demonstrated that the sRNA IsrR (originally named Tsr25 (24)) was required for growth during divalent metal starvation suggesting that it is a functional analog of RyhB or FsrA (25). The expression of IsrR resulted in decreased expression of Fe-dependent enzymes formate dehydrogenase (*fdhA*) and glutamate synthase (*gltB2*). A direct interaction was noted between IsrR and the *fdhA* transcript *in vitro*. The *isrR* locus was initially identified in a global transcriptomic analysis as the most upregulated sRNA in human serum (24). Consistent with a role in pathogenesis, an Δ*isrR* mutant caused decreased mortality in a murine septicemia model (25).

*S. aureus* relies on appropriate metabolic adaptation to host encountered stresses such as iron limitation (18). Additionally, central metabolism and TCA cycle function impacts virulence factor production and infection (26,27). This study was initiated to determine why a *S. aureus* Δ*fur* mutant has decreased expression of TCA cycle enzyme aconitase. We demonstrate that IsrR modulates the expression of *acnA* and directly interacts with the *acnA* mRNA transcript. We also demonstrate that IsrR alters the expression of additional genes coding TCA cycle enzymes. We also demonstrate that IsrR and Fur contribute to cellular iron homeostasis and virulence in murine infection models of skin and pneumonia.

## Materials and methods

### Chemicals, bacterial strains, and growth conditions

Unless specified, the *S. aureus* strains used in this study (Table 1) were isogenic and constructed in the community associated *S. aureus* MRSA strain USA300_LAC that was cured of the native plasmid pUSA03 that confers erythromycin resistance (28). All bacteria were grown at 37 °C in tryptic soy broth (TSB) (MP Biomedicals) or a chemically defined medium containing the 20 canonical amino acids with or without 10 mM glucose (3). Solid tryptic soy agar (TSA) and chemically defined media were generated by adding 1.5 % (w/v) agar (VWR). Liquid cultures were shaken at 200 rpm. TSB treated with Chelex 100 resin (Bio-Rad) was prepared as previously described (29). Unless stated otherwise, cells were cultured in 10 mL capacity culture tubes containing 2.0 mL of liquid medium. Liquid phenotypic analysis was conducted in 96-well microtiter plates containing 200 μL of media per well using a BioTek 808E visible absorption spectrophotometer with continuous shaking at shake speed high. The optical density of cultures was measured at 600 nm (A_600_). For quantitative growth, strains were grown overnight and washed with PBS before diluting to an optical density (A_600_) of 0.05 in 200 µL of media. For spot plate growth analyses using solid media, strains were cultured for 18 hours in TSB before harvesting by centrifugation. Cells were washed with PBS, standardized to an optical density of 2 (A_600_), serially diluted in PBS, and 5 μL aliquots were spotted upon solid media. Iron salts were added as ferrous sulfate.

**Table 1.** *Staphylococcus aureus* USA300_LAC strains used in this study.

| Strain name | Genotype | Reference |
| --- | --- | --- |
| JMB1100 | USA300_LAC wild type (WT) | A. Horswill |
| JMB10842 | $\Delta fur::tetM$ | (32) |
| JMB10495 | $\Delta fur::tetM$ *isrR* * | This study |
| JMB11112 | $\Delta fur::tetM$ SAUSA300_1309::Tn | This study |
| JMB11113 | $\Delta fur::tetM$ *isrR* * SAUSA300_1309::Tn | This study |
| JMB11292 | $\Delta isrR$ | This study |
| JMB11293 | $\Delta fur::tetM$ $\Delta isrR$ | This study |
| JMB1886 | *geh*::pLL39 | (61) |
| JMB11803 | *acnA*::Tn | (35) |
| JMB7868 | $\Delta acnA::tetM$ | (26) |
| JMB11804 | *acnA*::Tn $\Delta fur::tetM$ | This study |
| JMB11805 | *acnA*::Tn $\Delta isrR$ | This study |
| JMB11806 | *acnA*::Tn $\Delta fur::tetM$ $\Delta isrR$ | This study |
| JMB11448 | SAUSA300_1456::Tn *geh*::pLL39 | This study |
| JMB11449 | *fur*<sub>E11Stop</sub> SAUSA300_1456::Tn *geh*::pLL39 | This study |
| JMB11395 | $\Delta isrR$ SAUSA300_1456::Tn *geh*::pLL39 | This study |
| JMB11392 | *fur*<sub>E11Stop</sub> $\Delta isrR$ SAUSA300_1456::Tn *geh*::pLL39 | This study |
| JMB11393 | *fur*<sub>E11Stop</sub> $\Delta isrR$ SAUSA300_1456::Tn *geh*::pLL39 *isrR* | This study |
| JMB13983 | *fur*<sub>E11Stop</sub> $\Delta isrR$ SAUSA300_1456::Tn *geh*::pLL39 *isrR*_C1 | This study |
| JMB13984 | *fur*<sub>E11Stop</sub> $\Delta isrR$ SAUSA300_1456::Tn *geh*::pLL39 *isrR*_C2 | This study |
| JMB14314 | *fur*<sub>E11Stop</sub> $\Delta isrR$ SAUSA300_1456::Tn *geh*::pLL39 *isrR*_C1_C2 | This study |

Antibiotics were added at the following final concentrations: 100 μg mL^-1^ ampicillin (Amp); 10 μg mL^-1^ chloramphenicol (Cm) to select for plasmids and 3.3 μg mL^-1^ Cm to maintain plasmids (called TSB-Cm); 5 μg mL^-1^ erythromycin (Erm); 3 μg mL^-1^ tetracycline (Tet); 100 ng mL^-1^ anhydrotetracycline (Atet). Protein concentrations were determined using Bradford reagent (Bio-Rad Laboratories Inc., Hercules, CA). Unless stated otherwise, all chemicals were purchased from Sigma-Aldrich (St. Louis, MO).

### Plasmid and strain construction

The restriction minus strain *S. aureus* RN4220 was used for transformations (30) and transductions were done using bacteriophage 80α (31). 5α Competent *Escherichia coli* (NEB) cultured in lysogeny broth was used for plasmid preparation.

Synthetic DNA (Table S2) was synthesized by Twist Biosciences (San Francisco, CA) and DNA primers (Table S1) were synthesized by Integrated DNA Technologies (Coralville, IA). Plasmids are listed in Table 2. Quick Ligase, restriction enzymes, competent *E. coli*, and HiFi DNA Assembly kit were purchased from New England Biolabs (NEB). All bacterial strains were PCR or sequence verified before use. Plasmid DNA and PCR products were sequenced by Azenta Life Sciences (South Plainfield, NJ).

**Table 2.** Plasmids used in this study.

| **Name** | **Function** | **Reference** |
| --- | --- | --- |
| pJB38 | Mutant generation | (62) |
| pJB38_Δ*isrR* | Δ*isrR* mutant generation | This study |
| pEPSA5 | Inducible expression complementation | (63) |
| pEPSA5_*isrR* | Inducible expression *isrR* | This study |
| pEPSA5_as_*isrR* | Inducible expression *isrR* antisense | This study |
| pEPSA5_trunk_*isrR* | Inducible expression truncated *isrR* | This study |
| pEPSA5_ *isrR* * | Inducible expression *isrR* * | This study |
| pEPSA5_*acnA* | Non-native promoter expression | This study |
| pOS-1-*plgt* | Genetic complementation *lgt* promoter | (64) |
| pOS-1-*plgt_fur* | Genetic complementation *lgt* promoter | (5) |
| pOS_*pisrR_gfp* | *isrR* transcriptional reporter | This study |
| pOS_*pisrR\*_gfp* | *isrR* * transcriptional reporter | This study |
| pLL39 | Genetic complementation | (65) |
| pLL39_*isrR* | Native promoter complementation | This study |
| pLL39_*isrR_C1* | Native promoter complementation | This study |
| pLL39_*isrR_C2* | Native promoter complementation | This study |
| pLL39_*isrR_C1_C2* | Native promoter complementation | This study |
| pOS_*plgt_acnA_gfp* | *acnA* translational reporter | This study |
| pOS_*plgt_acnA1_gfp* | Mutated *acnA* translational reporter | This study |

The pJB38_Δ*isrR* was constructed by combining two digested PCR amplicons corresponding to the DNA upstream and downstream of the *isrR* locus with digested pJB38. The upstream region of *isrR* was amplified using primer pair tsr25 AF and tsr25 AR. The downstream region was amplified using primer pair tsr25 BF and tsr25 BR. pJB38 was digested using SacI and KpnI. The upstream amplicon was digested with SacI and MulI. The downstream region was digested with MulI and KpnI.

The pEPSA5_*isrR* complementation vector was created using the tsr25 pEPSA 5 EcoRI and tsr25 pEPSA 3 SalI primer pair with JMB1100 as template DNA. The pEPSA5_as_*isrR* complementation vector was created using the tsr25 pMAL 5 SalI and tsr25 pMAL 3 EcoRI primer pair with JMB1100 as template DNA. The pEPSA5_*isrR*\* vector was created using the tsr25* pEPSA 5 EcoRI and tsr25 pEPSA 3 SalI primer pair with JMB10495 cells as template DNA. The pEPSA5_trunk_*isrR* was created using the tsr25 trunk pEPSA 5 EcoRI and tsr25 pEPSA 3 SalI primer pair with JMB1100 cells as the DNA template. The pEPSA5_*acnA* expression vector was created using the pEPSA5_acnAEcoRI (acnA RBS) and pEPSA_acnA3SalI primer pair. All pEPSA5 vectors were digested with EcoRI and SalI.

The pLL39_*isrR* complementing plasmid, pLL39_*isrR_C1*, pLL39_*isrR*_C2, and pLL39_*isrR_C1_C2* were created using the Tsr25 comp 5 SalI and Tsr25 comp 3 BamHI primer pair with JMB1100, synthetic DNA *isrR_C1,* synthetic DNA *isrR*_C2, and synthetic DNA *isrR_C1_C2* used as template DNA, respectively. All pLL39 vectors were digested with SalI and BamHI.

The pOS_p*isrR*_*gfp* and pOS_p*isrR*\*_*gfp* transcriptional reporters were constructed using the pOStsr25_2forHindIII and pOStsr25_2revkpnI primer pair with JMB1100 and JMB10495 cells as template DNA, respectively. PCR products were digested and ligated into KpnI and HindIII digested pOS_*saeP1_gfp*.

The pOS_p*lgt_acnA_gfp* and pOS_p*lgt_acnA*1*_gfp* translational reporters were generated using pOS_p*lgt* digested with NdeI. The aconitase insert of pOS_p*lgt_acnA_gfp* was created using the plgt_acnA and acnA_gfp Wt A rev primer pairs with JMB1100 as template DNA. The *gfp* insert of pOS_p*lgt_acnA_gfp* was created using the acnA_gfp wt A for and gfp_plgt rev primer pair with the pOS_*saeP1_gfp* plasmid as template DNA. The aconitase insert of pOS_p*lgt_acnA1_gfp* was created using the plgt_acnA and acnA_gfp MutC rev primer pair with JMB1100 as template DNA. The *gfp* insert of pOS_p*lgt_acnA1_gfp* was created using the acnA_gfp MutC for and gfp_plgt rev primer pair with the pOS_*saeP1_gfp* plasmid as template DNA. All digested pOS_p*lgt* vectors, *acnA*, and *gfp* inserts were ligated using the NEB Hifi DNA Assembly Master Mix.

### RNA extraction, cDNA synthesis and qPCR

To analyze RNA abundances corresponding to *isrR*, *S. aureus* strains were cultured overnight and diluted into 2 mL TSB to an optical density (A_600_) of 0.1 in 10 mL culture tubes. The cell cultures were incubated with shaking until an A_600_ of 0.5 before 1 mL of cells was harvested by centrifugation, washed with PBS, and resuspended in 500 µL RNAprotect (QIAGEN).

To analyze *isrR* transcript stability, the Δ*fur::tetM* Δ*isrR* strain containing pEPSA5_*isrR* or pEPSA5_*isrR*\* were cultured overnight in TSB with 10 μg mL^-1^ chloramphenicol in 10 mL culture tubes at 37 °C with shaking. Cultures were diluted into 5 mL of fresh TSB with 10 μg mL^-1^ chloramphenicol and 2% xylose to an A_600_ of 0.1 in 30 mL culture tubes. The cells were cultured with shaking until an A_600_ of 0.5. Next, 1.5 mL of culture were added to 10 mL culture tubes containing rifampicin (100 µg mL^-1^) and incubated with shaking at 37 °C. Cells were harvested at indicated time points by centrifugation, washed with PBS, and resuspended in 500 µL RNAprotect (QIAGEN). RNA extraction, cDNA synthesis, and transcript quantification (QuantStudio 3, Bio-Rad Laboratories Inc., Hercules, CA) were performed as previously described (29).

### Northern blot analyses

RNA (3 μg/lane) was loaded onto a formaldehyde agarose gel and electrophoresed for 1 h 30 min at 120 V. RNAs were transferred onto a positively charged nylon membrane overnight by capillary transfer with 20 x SSC (1x SCC is 0.15 M NaCl plus 0.015 M sodium citrate) buffer and UV-crosslinked to the membrane. The presence of rRNA and ladder bands were visualized by staining the membrane with methylene blue (0.04 % in 0.5 M acetate solution). To detect *isrR*, a radio-labeled probe was made as follows: PCR targeting *isrR* using primers isrR5north and isrR3north was performed and the PCR mixture was radiolabeled using the Roche random prime labeling kit. Approximately 1 μg of PCR product was used with [α-32P] ATP according to the manufacturer’s protocol. Probes were purified using Illustra MicroSpin G-25 columns (GE Healthcare). Membranes were prehybridized overnight at 45 °C in ULTRAhyb-Oligo buffer (Thermo Scientific) and then incubated with radiolabeled probe overnight at 45°C. After incubation, membranes were washed with 2×, 1×, and 0.5× SSC buffer and visualized using a phosphor imager screen.

### Suppressor screen, genome sequencing, and SNP mapping

Ten independent cultures of the Δ*fur::tetM* strain were grown overnight in 2 mL of TSB in 10 mL capacity culture tubes. One mL of cells was pelleted by centrifugation, resuspended in 1 mL PBS, and then diluted 1:100 in PBS. One hundred µL of the dilution was spread on chemically defined agar media containing amino acids without glucose. One colony from each plate (i.e. one per culture for ten total) was retained and struck for isolation. The increased growth phenotypes were verified by serially diluting cultures and spot plating them on solid defined amino acid media with and without glucose. We repeated the assay for a second time with five independent overnight cultures for a total of 15 suppressed strains.

For chromosomal DNA isolation, the Δ*fur::tetM* suppressed strains and the Δ*fur::tetM* (JMB10842) parent strain were cultured overnight in 2 mL of TSB and genomic DNA was purified using the MaterPure Gram positive genomic DNA purification kit (LGC Biosearch Technologies). DNA was sequenced by SeqCenter (Pittsburgh, PA, USA) using Illumina technology.

Whole genome sequencing data was analyzed using the CLC Genomics Workbench software package (Qiagen). Reads were aligned to the *S. aureus* genome, using the USA300_FPR3757 genome sequence as a reference as previously described (32). Quality-based variant detection was then performed to identify polymorphisms in each strain. A minimum threshold detection frequency of 80% was employed. The lists of polymorphisms generated for each suppressor mutant strain were cross-referenced against the parental strain (JMB10842). Common polymorphisms were eliminated (as were polymorphisms in homopolymeric nucleotide tracts) resulting in the identification of specific genetic variations between the suppressor strains and parental strain. The SNP in the promoter of the *isrR* gene was confirmed in the suppressed strains using Sanger sequencing. These strains were referred to as Δ*fur::tetM isrR\** and JMB10495 was used as a representative strain.

### Transcriptional reporter assays

Overnight cultures of *S. aureus* strains containing a plasmid-based transcriptional reporter were cultured overnight in 2 mL TSB supplemented with 3.3 μg mL^-1^ chloramphenicol in 10 mL culture tubes at 37 °C with shaking. Cultures were then diluted to an optical density (A_600_) of 0.05 in triplicate into 2 mL TSB supplemented with 3.3 μg mL^-1^ chloramphenicol +/- 250µM DIP in 10 mL culture tubes and incubated at 37 °C with shaking for 16 hours. Optical density (A_600_) and GFP fluorescence (excitation 485 nm, emission 520 nm) were measured in microtiter plates using a Varioskan Lux plate reader (Thermo Scientific).

### EMSA assays

PCR fragments containing T7-*acnA* (from -41 to +541), T7-*citM* (from -46 to +725), T7-*citZ* (from -41 to +850), T7-*lukH* (from -100 to +807), T7-*mqo* (from -248 to +328), and T7-*sdhC* (full-length, from -260 to +615) were used as DNA template for in vitro transcription with T7 RNA polymerase. RNAs were finally purified and radiolabeled when required as previously described (33).

5’-radiolabelled IsrR (20,000 cpm/sample, concentration <1 pM) and above-mentioned cold RNAs were separately denatured at 90°C in the buffer GR-(20 mM Tris-HCl pH 7.5, 60 mM KCl, 40 mM NH_4_Cl, 3 mM DTT), cooled 1 min on ice, and incubated at room temperature for 15 min in presence of 10 mM MgCl_2_. Renatured RNAs were then mix and incubated at 37°C for 15 min. Finally, samples were loaded on a 6% polyacrylamide gel under non-denaturing conditions (300 V, 4°C). Results are representative of two independent experiments.

### Enzyme assays

#### Aconitase assays

Aconitase (AcnA) assays were conducted as previously described (34). Briefly, strains were cultured overnight in 2 mL of TSB with or without 10 μg mL^-1^ Cm before diluting them to an optical density (A_600_) of 0.05 in 2 mL of TSB in 10 mL culture tubes . For strains containing pEPSA5, strains were cultured in TSB supplemented with 0.25 % or 2 % xylose for pEPSA_*acnA* and pEPSA_*isrR* containing strains respectively, and 3.3 μg mL^-1^ Cm. Strains were cultured with shaking at 200 rpm at a 45 degree angle for 16 hours. After incubation, one mL of cells was pelleted down and washed twice with PBS. After assaying AcnA activity as previously described (35), protein concentrations were determined using Bradford protein colorimetric assay modified for 96-well plate (Bio-Rad Protein Assay Dye Reagent Concentrate).

#### Succinate dehydrogenase assays

Succinate dehydrogenase (Sdh) assays were conducted as previously described (36). Cells were cultured and lysed as described for AcnA assays. The oxidation of succinate by succinate dehydrogenase was followed spectrophotometrically using the redox dye 2,6-dichlorophenol indophenol. The reaction mixture (1 mL) contained: potassium phosphate buffer (0.1 M, pH 7.4), KCN (3 mM), 2,6-dichlorophenol indophenol (DCPIP, 25 µM), *N*-methylphenazonium methosulphate (2.2 mM), succinic acid (20 mM, pH 7) and 20 µL of cell lysate. The reduction of DCPIP (molar extinction coefficient 21 mM^-1^ cm^-1^) was followed at 600 nm for two minutes after addition of all reagents as described (37). The succinate dependent slope (difference between the slope of absorbance/time of the sample with succinate minus the slope of the sample without succinate) was used to calculate specific Sdh activity. Protein concentrations were determined as described for the aconitase assay.

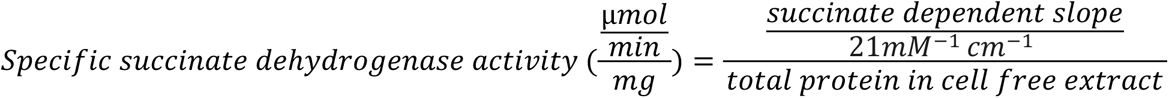

#### Malate quinone oxidoreductase assays

Cells were cultured and lysed as described for AcnA assays. Reaction mixture was the same as described for Sdh assays except for the use of sodium malate (20 mM) instead of succinic acid.

### Streptonigrin sensitivity

Cells were cultured overnight and diluted to an optical density (A_600_) of 0.05, after which 100 µL of the dilute culture was added to 4 mL of soft TSB agar (0.3 % wt vol^-1^) and overlaid on TSA media. After soft agar solidified, 2.5 µL of streptonigrin (1 mg mL^-1^) was spotted. Zone of clearance was measured after one day incubation at 37°C.

### CAS siderophore assay

Overnight cultures in TSB were diluted 100-fold into 1 mL of Chelex (Bio-Rad)-treated TSB with the addition of 25 μM zinc acetate, 25 μM MnCl_2_, 1 mM MgCl_2_ and 100 μM CaCl_2_ in 10 mL glass culture tubes. The cultures were incubated at 37 °C with shaking for 18 hours. The chrome azurol S siderophore assay was performed on the spent media using the modified microplate method as previously reported (38,39).

### Whole cell metal quantification

*S. aureus* strains were grown for 18 hours overnight in TSB before diluting them to an optical density (A_600_) of 0.05 into 7.5 mL of TSB or Chelex-treated (Bio-Rad) TSB in 30 mL capacity culture tubes as described previously (29). Cells were allowed to grow with shaking for eight hours. Pre-weighted metal-free 15 mL propylene tubes were used to pellet the cells in a tabletop centrifuge at 4 °C (Eppendorf, Hauppauge, NY). Pellets were washed three times with 10 mL of ice-cold PBS. Samples were kept at -80 °C or on dry ice until processing.

Cell pellets were acid digested with 2 mL of Optima grade nitric acid (ThermoFisher, Waltham, MA) and 500 μL hydrogen peroxide (Sigma, St. Louis, MO) for 24 hr at 60 °C. After digestion, 10 mL of UltraPure water (Invitrogen, Carlsbad, CA) was added to each sample. Elemental quantification on acid-digested liquid samples was performed using an Agilent 7700 inductively coupled plasma mass spectrometer (Agilent, Santa Clara, CA). The following settings were fixed for the analysis Cell Entrance = −40 V, Cell Exit = −60 V, Plate Bias = −60 V, OctP Bias = −18 V, and collision cell Helium Flow = 4.5 mL min^−1^. Optimal voltages for Extract 2, Omega Bias, Omega Lens, OctP RF, and Deflect were determined empirically before each sample set was analyzed. Element calibration curves were generated using ARISTAR ICP Standard Mix (VWR). Samples were introduced by a peristaltic pump with 0.5 mm internal diameter tubing through a MicroMist borosilicate glass nebulizer (Agilent). Samples were initially up taken at 0.5 rps for 30 s followed by 30 s at 0.1 rps to stabilize the signal. Samples were analyzed in Spectrum mode at 0.1 rps collecting three points across each peak and performing three replicates of 100 sweeps for each element analyzed. Sampling probe and tubing were rinsed for 20 s at 0.5 rps with 2% nitric acid between each sample. Data were acquired and analyzed using the Agilent Mass Hunter Workstation Software version A.01.02.

### Murine models of infection

#### Acute pneumonia model

Experiments were conducted as previously described (40). Briefly, mice were intranasally infected with 2-4 x 10^7^ colony forming units (CFU) in 50 μL of PBS under anesthesia (ketamine and xylazine). Bacterial loads were enumerated at 24 h post infection from bronchoalveolar lavage fluid (BALF) by washing the airway 3 times with 1 mL of PBS, and homogenized lung tissue. Bacterial counts were quantified by serial dilution using CHROMagar *S. aureus* plates (BD Biosciences).

#### Skin and soft tissue model

All studies were conducted in accordance with an approved protocol at The University of Kansas Medical Center. Female C57BL/6J mice (Jackson Laboratories) were used in a subcutaneous skin infection model as previously described (41). Briefly, 8-week-old mice were injected with ∼2.1e7 CFU of mid-exponential phase cells suspended in PBS. Mice were imaged daily, and lesion sizes determined using ImageJ. On the final day, mice were euthanized and the lesion plus ∼3 mm of neighboring tissue were homogenized in 1X HBSS 0.2% HAS 10 mM HEPES buffer using an MP Biomedicals Fastprep-24 homogenizer with Lysing Matrix H tubes following the manufacture’s protocol for skin. A sample was taken to determine bacterial titers by dilution plating. Next, the samples were clarified by centrifugation, treated with 1x protease inhibitor (Roche), and frozen at -80 °C until processed for cytokines. Cytokines were measured using the BD cytometric bead array (CBA) mouse flex set protocol on a BD Aria flow cytometry machine and analyzed in FCS express.

### Ethics statement

Animal work in this study was carried out in strict accordance with the recommendations in the Guide for the Care and Use of Laboratory Animals of the NIH (National Academies Press, 2011), the Animal Welfare Act, and US federal law. Protocols were approved by the Institutional Animal Care and Use Committee of Rutgers New Jersey Medical School of Newark, New Jersey, USA, as well as The University of Kansas Medical Center, Kansas City, Kansas, USA.

### Statistical analysis

For two group comparisons (controls vs treatment or between bacterial strains), student’s t-tests were performed. Multiple group comparisons for animal data were performed using an ANOVA with a Kruskal-Wallis test or a Mann-Whitney non-parametric test for two group comparisons. All analyses were conducted with Sigmaplot 11, Microsoft Excel, or Prism 9.

## Results

### Iron starvation decreases aconitase activity in a Fur-dependent manner

Previous work demonstrated that both the absence of Fur in *S. aureus* or Fe depletion resulted in a redirection of central metabolism where glycolysis is increased, and the TCA cycle is downregulated (22,23). The shift towards fermentative metabolism increases Fe availability, and since the TCA cycle contains Fe-S cluster requiring enzymes, its downregulation could be a way to decrease nonessential Fe usage in an Fe sparing response (42).

We tested the hypothesis that the Fe-S cluster requiring enzyme aconitase (AcnA), which converts citrate to isocitrate in the TCA cycle, would have decreased activity upon Fe limitation. We quantified the activity of AcnA in the parent strain, USA300_LAC, and the isogenic Δ*fur::tetM* mutant. The strain lacking Fur had nearly undetectable AcnA activity and the phenotype could be genetically complemented (Fig 1A). The divalent metal chelator 2,2’-dipyridyl (DIP) has a high affinity for Fe(II) and co-culture with DIP results in Fur derepression (42,43). When the WT strain was co-cultured with DIP the activity of AcnA was significantly decreased (Fig 1B).

**Figure 1.**
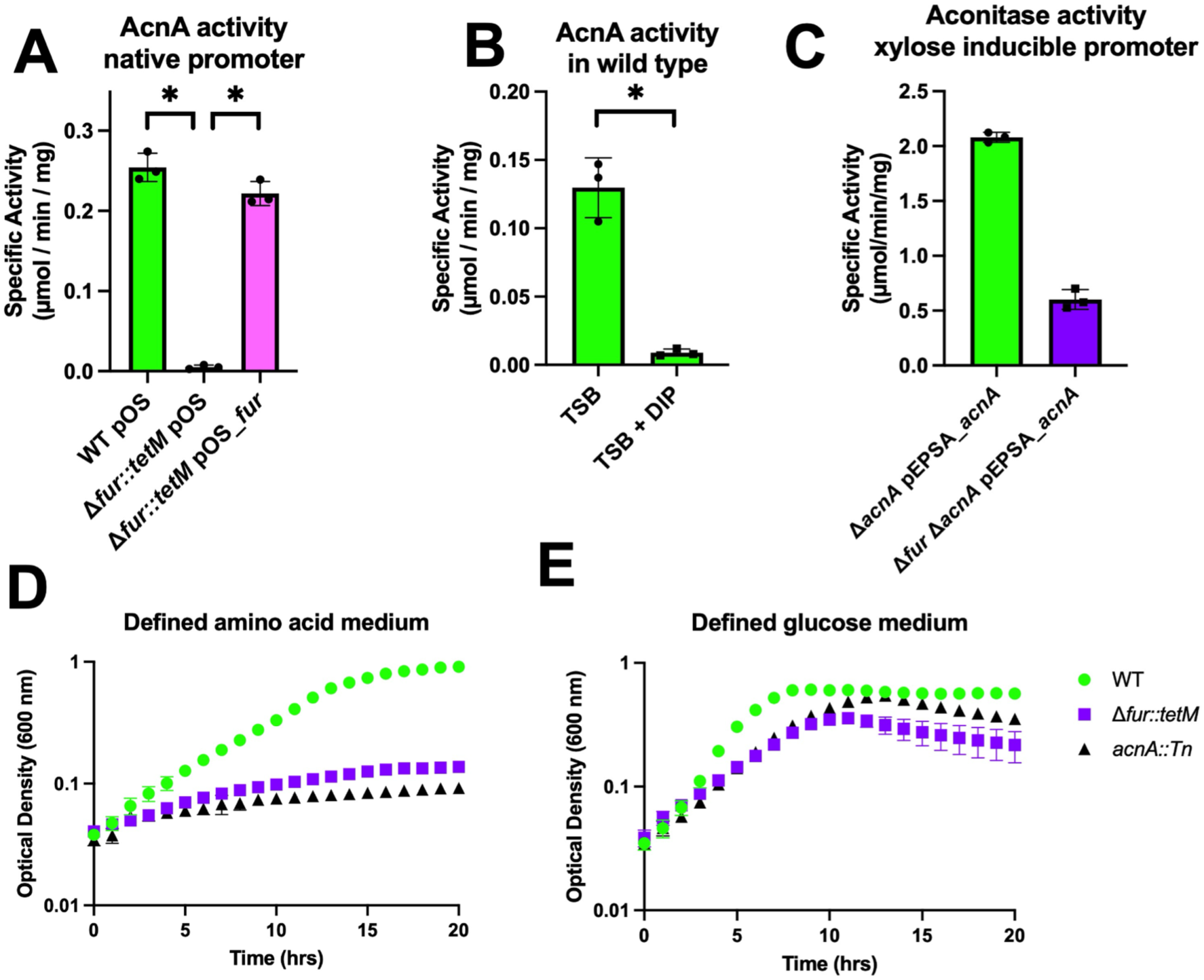
A *S. aureus* Δ*fur* mutant has decreased aconitase expression. **Panel A.** Aconitase activity was quantified in cell free lysates harvested from the wild type (WT) (JMB1100) and Δ*fur::tetM* (JMB10842) strains carrying pOS-1-p*lgt* or pOS-1-p*lgt_fur* after culture in TSB-Cm medium. **Panel B.** Aconitase activity was quantified in cell free lysates from the WT after culture in TSB +/- 250 µM 2,2’ dipyridyl (DIP). **Panel C.** Aconitase activity was quantified in cell free lysates harvested from the *acnA::Tn* (JMB11803) and Δ*fur::tetM acnA::Tn* (JMB11804) strains with pEPSA5_*acnA* after culture in TSB-Cm medium supplemented with 0.25% xylose. **Panel D.** Culture optical densities (A_600_) were monitored for the WT, Δ*fur::tetM,* and *acnA::Tn* strains in a defined medium containing amino acids as the sole carbon and energy sources. **Panel E.** Culture optical densities (A_600_) of the WT, Δ*fur::tetM,* and *acnA::Tn* strains were monitored in a liquid defined medium containing amino acids supplemented with 10 mM glucose. The data shown represent the average of biological triplicates with standard deviations shown. Error bars are shown for all data but in some cases (Panels D and E) are smaller than the symbols used. Student’s two-tailed t-tests were performed on the data and * represents a p-value of <0.05.

We next examined whether decoupling *acnA* transcriptional activity from its native promoter would increase AcnA activity in the *fur* mutant. To this end, we placed *acnA* and its ribosomal binding site (RBS) under the transcriptional control of a xylose inducible promoter (*xylRO*) using the pEPSA5 plasmid. We examined AcnA activity in the extracts of the *acnA::Tn* and *acnA::Tn* Δ*fur::tetM* strains containing pEPSA5_*acnA* after culture with xylose. The *acnA::Tn* Δ*fur::tetM* strain had significantly decreased AcnA activity compared to the *acnA::Tn* strain (Fig 1C). These data demonstrate that *acnA* expression is altered in a strain lacking *fur* and this regulation is also independent of its native promoter.

Using amino acids for carbon and energy in *S. aureus* requires a functional TCA cycle (34,44). We compared growth of the WT, Δ*fur::tetM*, and *acnA::Tn* strains cultured in a chemically defined medium containing amino acids with and without glucose. The Δ*fur::tetM* and *acnA::Tn* strains had severe growth defects in the medium containing amino acids as the sole carbon and energy source whereas the WT was capable of growth (Fig 1D). All three strains were capable of growth when the medium was supplemented with glucose (Fig 1E). We also found that the Δ*fur::tetM* and *acnA::Tn* strains did not grow on solid chemically defined medium containing amino acids for carbon and energy. Again, this phenotype was reversed by supplementing the medium with glucose (Fig S1).

### A null mutation in *isrR* promotes growth of a Δ*fur* strain on amino acids

We conducted suppressor analysis to provide insight into the defective growth of the Δ*fur::tetM* strain on an amino acid medium. We individually plated fifteen cultures of the Δ*fur::tetM* strain on a chemically defined solid medium containing only amino acids as carbon and energy sources. We isolated one colony from each plate, verified the phenotype of improved growth with amino acids, and then determined the locations of the single nucleotide polymorphisms by whole genome sequencing. All fifteen strains contained a C→T mutation located 604 base pairs upstream of the translation start site of *arlR* (SAUSA300_1308). This chromosomal location corresponds to *tsr25* (recently renamed *isrR*), which codes for a sRNA (24,25). The mutations correspond to a G→A change within the *isrR* sequence at the +1 location as determined by RNA sequencing (45) and the +3 location as determined by 5’ RACE analysis (25). Henceforth, we refer to this allele as *isrR*\*. The *isrR\** mutation permitted growth of the Δ*fur::tetM* mutant on both solid and liquid chemically defined amino acid media (Fig 2A and Fig S2). The Δ*fur::tetM isrR\** strain also had increased AcnA activity compared to the Δ*fur::tetM* mutant (Fig 2B). Three findings suggested that the *isrR*\* allele decreased IsrR function, and thereby corrected the phenotypes of the Δ*fur::tetM* strain. First, we linked a transposon (Tn) (SAUSA300_1309::Tn) to the *isrR*\* mutation and then used this strain as a DNA donor for transduction into the Δ*fur::tetM* strain. We isolated two classes of transductants. One class corrected the growth of the Δ*fur::tetM* mutant on amino acid media and the second class did not. Sanger sequencing determined that the strains that grew on amino acid medium contained the *isrR*\* allele, and all strains that did not grow contained the wild type *isrR* allele. Second, we constructed Δ*isrR* and Δ*fur::tetM* Δ*isrR* strains. Deletion of *isrR* in the Δ*fur::tetM* strain permitted growth on solid and liquid defined amino acid medium (Fig 2C and Fig S3). Deletion of *isrR* in the Δ*fur::tetM* strain also increased AcnA activity (Fig 2D). Introduction of the *acnA::Tn* mutation into the Δ*fur::tetM* Δ*isrR* strain prevented growth in amino acid medium consistent with the hypothesis that the increased growth imparted by the null *isrR* mutations requires functional AcnA (Fig 2C and Fig S3). Third, we genetically complemented the Δ*isrR* strains. The pLL39 episome codes for tetracycline resistance so we could not use the Δ*fur::tetM* strain. Instead, we used previously described strains containing a null *fur*_E11stop_ allele (*fur*\*) that is genetically linked to a transposon in gene SAUSA300_1456 (42). As expected, the strain containing *fur*\* had greatly reduced AcnA activity. The presence of the Δ*isrR* mutation in the *fur*\* strain increased AcnA activity and growth on chemically defined amino acid medium and the phenotypes could be genetically complemented (Figs 2E and S4).

**Figure 2.**
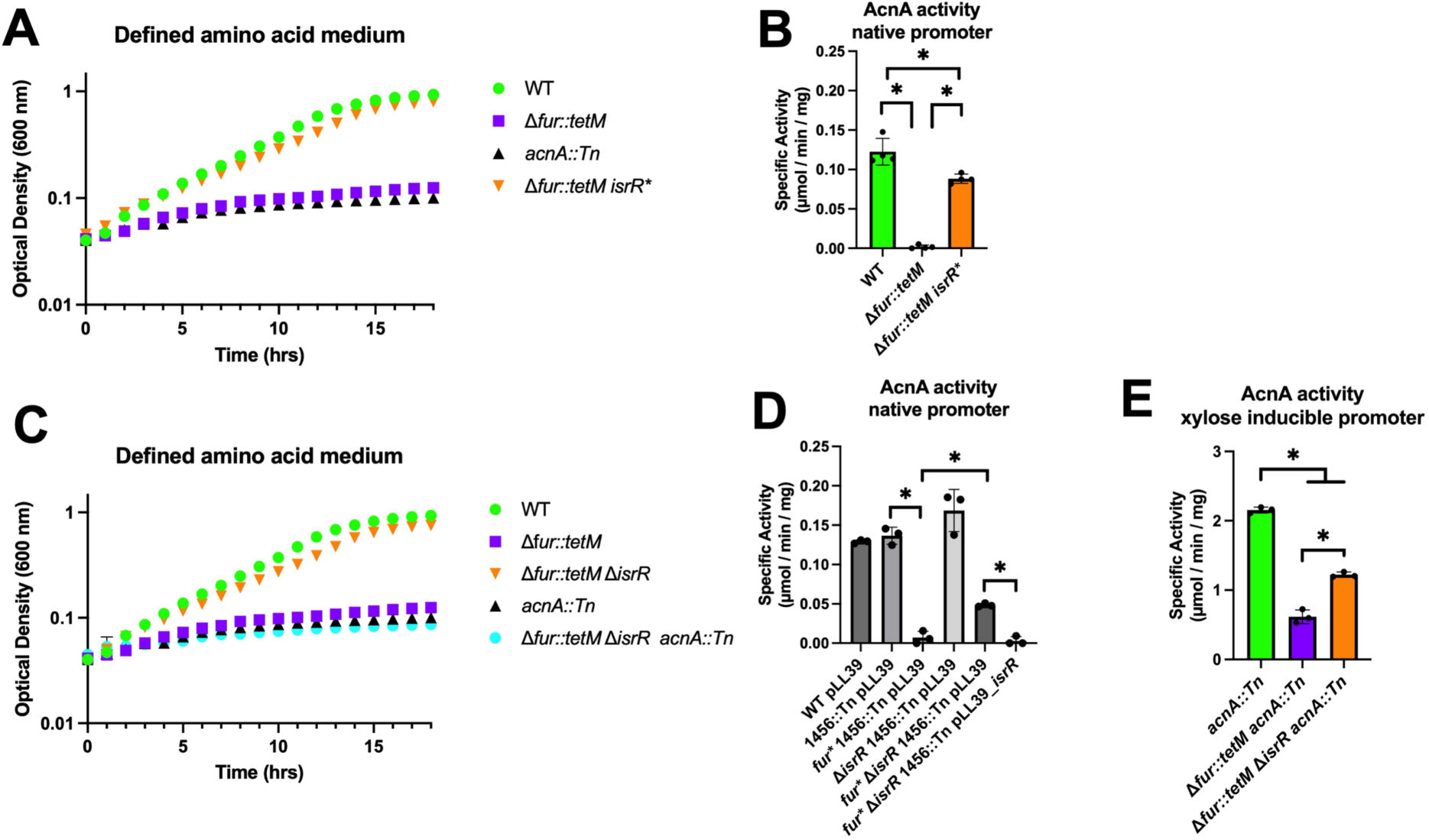
A null mutation in *isrR* suppresses the amino acid growth defect and increases aconitase activity of a Δ*fur* mutant. **Panel A.** Culture optical densities (A_600_) of the wild type (WT) (JMB1100), Δ*fur::tetM* (JMB10842), Δ*fur::tetM isrR\** (JMB10495), and *acnA::Tn* (JMB11803) strains were monitored when cultured on defined medium containing amino acids as carbon and energy sources. **Panel B.** Aconitase activity was quantified in cell free lysates harvested from the WT, Δ*fur::tetM,* and Δ*fur::tetM isrR\** strains after culture in TSB medium. **Panel C.** Culture optical densities (A_600_) of the WT, *acnA::Tn,* Δ*fur::tetM,* Δ*fur::tetM* Δ*isrR* (JMB11293), and Δ*fur::tetM* Δ*isrR acnA::Tn* (JMB11806) strains were monitored in liquid defined medium containing amino acids as carbon and energy sources. **Panel D.** Aconitase activity was quantified in cell free lysates harvested from the WT with pLL39 (JMB1886), SAUSA300_1456::Tn (*1456::Tn*) pLL39 (JMB11448), *fur\** 1456::Tn pLL39 (JMB11449), Δ*isrR* 1456::Tn pLL39 (JMB11395)*, fur\** Δ*isrR 1456::Tn* pLL39 (JMB11392), and *fur\** Δ*isrR 1456::Tn* pLL39_*isrR* (JMB11393) strains after culture in TSB medium. **E.** Aconitase activity was quantified in cell free lysates harvested from the *acnA::Tn,* Δ*fur::tetM acnA::Tn* (JMB11804), and Δ*fur::tetM* Δ*isrR acnA::Tn* strains containing pEPSA5_*acnA* after culture in TSB-Cm medium supplemented with 0.25% xylose. The data shown represent the average of biological triplicates with standard deviations shown. Error bars are shown for all data but in some cases (Panels D and E) are smaller than the symbols used. Student’s two-tailed t-tests were performed on the data and * represents a p-value of <0.05.

We also examined the effect of the Δ*isrR* mutation on AcnA activity when *acnA* was expressed from a non-native promoter. We quantified AcnA activity in cell lysates generated from the *acnA::Tn*, *acnA::Tn* Δ*fur::tetM*, and *acnA::Tn* Δ*isrR* Δ*fur::tetM* strains containing pEPSA5_*acnA* after culture in tryptic soy broth (TSB) containing xylose (Fig 2E). Again, the activity of AcnA was decreased in the strain lacking Fur introduction of the Δ*isrR* mutation abolished this decrease.

### The *isrR*\* allele decreases *isrR* transcription

The *isrR* promoter contains two near consensus Fur box sequences (25). The *isrR*\* mutation resulted in a base change in the Fur box proximal to the *isrR* transcription start site (Fig 3A). We conducted Northern blot analyses to 1) determine if *isrR* transcript abundances responded to divalent metal starvation, and 2) verify that it was regulated by Fur in USA300_LAC. The *isrR* transcript increased in abundance as we increased the concentration of DIP in the growth medium (Fig 3B). Moreover, the *isrR* transcript accumulated in the Δ*fur::tetM* strain when compared to the unchallenged WT strain, but there was not additional accumulation in the Δ*fur::tetM* strain upon co-culture with DIP (Fig 3C). These data are consistent with previous results demonstrating that *isrR* transcription is regulated by Fur (25) and suggest that Fur is the dominant divalent metal-dependent transcriptional regulator controlling *isrR* transcription.

**Figure 3.**
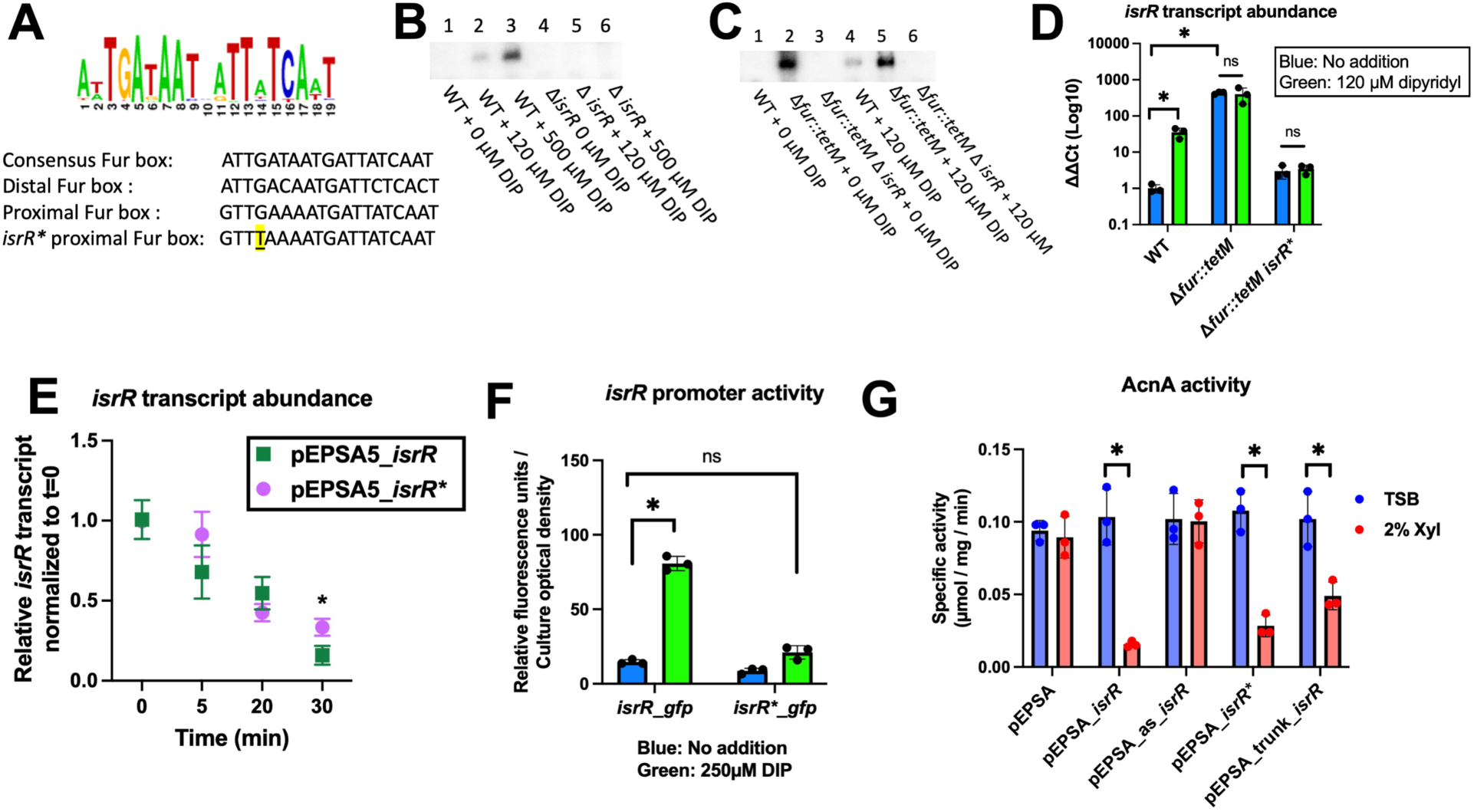
The *isrR\** mutation suppresses the phenotypes of the Δ*fur* mutant by decreasing *isrR* transcription. **Panel A.** Top, predicted consensus Fur box sequence as determined by RegPrecise, and bottom, consensus Staphylococcal Fur box, *isrR* distal Fur box, *isrR* proximal Fur box, and proximal Fur box in the *isrR\** strains with change highlighted and underlined. **Panel B.** Northern blot analysis of *isrR* transcripts using total RNA isolated from wild type (WT)(JMB1100) and Δ*isrR* (JMB11292) strains after culture in TSB media containing 0, 120, or 500 µM 2,2’ dipyridyl (DIP). **Panel C.** Northern blot analysis of *isrR* transcripts using total RNA isolated from WT, Δ*fur::tetM* (JMB10842), and Δ*fur::tetM* Δ*isrR* (JMB11293) strains after culture in TSB media with or without 120 µM DIP. **Panel D.** IsrR transcript abundance in the WT, Δ*fur::tetM,* and Δ*fur::tetM isrR\** strains after culture in liquid TSB media supplemented with or without 120 µM DIP. Transcript abundance was determined by quantitative PCR. **Panel E.** Quantification of transcripts corresponding to *isrR* after the Δ*fur::tetM* Δ*isrR* strain carrying either pEPSA5_*isrR* or pEPSA5_*isrR\** were cultured in TSB medium containing 2% xylose and subsequently rifampicin was added (t=0) to inhibit transcription. Transcript abundances were normalized to t=0. **Panel F.** Relative fluorescence of the WT strain containing the pOS_p*isrR_gfp* or pOS_p*isrR*_gfp* transcriptional reporter after culture in TSB-Cm with or without 250 µM DIP. **Panel G.** Aconitase activity in cell free lysates from the WT strain carrying pEPSA5, pEPSA5_*isrR*, pEPSA5_as_*isrR,* pEPSA5_*isrR\**, and pEPSA5_trunk_*isrR* after culture in TSB-Cm with or without 2% xylose. Panels B and C contain representative Northern blots. The data displayed in panels D-G represent the average of biological triplicates with standard deviations shown. Student’s two-tailed t-tests were performed on the data and * represents a p-value of <0.05.

To evaluate how the *isrR\** mutation was affecting the expression of *isrR*, we used quantitative PCR to determine the abundance of the transcripts corresponding to *isrR* in the WT, Δ*fur::tetM*, and Δ*fur::tetM isrR\** strains after culture in the presence and absence of DIP. The abundance of the *isrR* transcript was increased in the WT upon co-culture with DIP (Fig 3D). The *isrR* transcript was elevated in the Δ*fur::tetM* strain when compared to the WT and it did not further increase upon co-culture with DIP, which is consistent with the Northern analyses. The abundances of *isrR* transcripts in the Δ*fur isrR\** strain was like that of the WT. However, the *isrR* transcript abundance did not increase in the Δ*fur isrR\** strain upon co-culture with DIP. These data suggest that the *isrR*\* mutation decreases transcriptional activity of the *isrR* locus or that it promotes decreased IsrR transcript stability.

We tested the hypothesis that the *isrR*\* mutation was suppressing the defects of the Δ*fur* strain by decreasing IsrR stability. We cultured the Δ*fur::tetM* Δ*isrR* strain containing pEPSA5_*isrR* or pEPSA5_*isrR*\* in the presence of xylose to induce *isrR* transcription. We then added rifampicin to inhibit RNA synthesis, isolated samples at different time points, and quantified *isrR* transcripts. The transcripts corresponding to *isrR* and *isrR\** decayed at similar rates. In fact, the *isrR\** transcript was slightly more abundant than the *isrR* transcript 30 minutes after transcription was halted (Fig 3E).

We next tested the hypothesis that the *isrR*\* mutation was suppressing the defects of the Δ*fur* strain due to decreased *isrR* transcription. We generated constructs where the *isrR* or *isrR*\* promoter drove transcription of *gfp*. The WT strain containing either construct was cultured in TSB with and without DIP and *gfp* fluorescence was quantified. Co-culture with DIP significantly increased *gfp* expression in the strain containing the *isrR_gfp* transcriptional reporter (Fig 3F). However, the strain containing the *isrR*_gfp* did not display increased *gfp* expression with DIP.

We individually placed the expression of *isrR*, *isrR\**, antisense *isrR* (as_*isrR*), and a truncated *isrR* that begins 2 nucleotides downstream from the nucleotide mutated in *isrR\** (trunk_*isrR*), under the transcriptional control of *xylRO* promoter using pEPSA5. Induced expression of *isrR*, *isrR\**, or trunk_*isrR* in the WT strain significantly decreased AcnA activity and decreased growth on solid defined media compared to the WT strain containing the empty vector (Figs 3G and S4). The AcnA activity or the growth of WT expressing the as_*isrR* was not significantly different to that of the WT containing empty vector. These results suggest that 1) expression of *isrR* under metal replete conditions decreases *acnA* expression, and 2) the *isrR\** mutation does not affect the ability of the *isrR\** transcript to decrease *acnA* expression. Taken together, these findings are consistent with the hypothesis that the *isrR*\* mutation results in decreased transcription of the *isrR* locus, and thereby, suppresses the phenotypes of the Δ*fur::tetM* strain.

### The presence of IsrR results in decreased *acnA* expression

Our results are consistent with a model wherein the absence of Fur or upon Fe ion limitation, *isrR* is expressed and mediates the repression of *acnA* expression. We next tested the hypothesis that IsrR alters *acnA* expression post-transcriptionally. We used the program IntaRNA (Freiburg RNA tools (46,47)) to help predict potential interactions between IsrR and the *acnA* transcript. One potential interaction site was identified that overlayed the 5’ untranslated region and the first two bases of coding sequence with a base-pair minimal annealing energy of -14.2 kcal mol^-1^. This interaction included a 100% overlap with the Shine Dalgarno sequence (AGGGGG) (Figure 4A).

**Figure 4.**
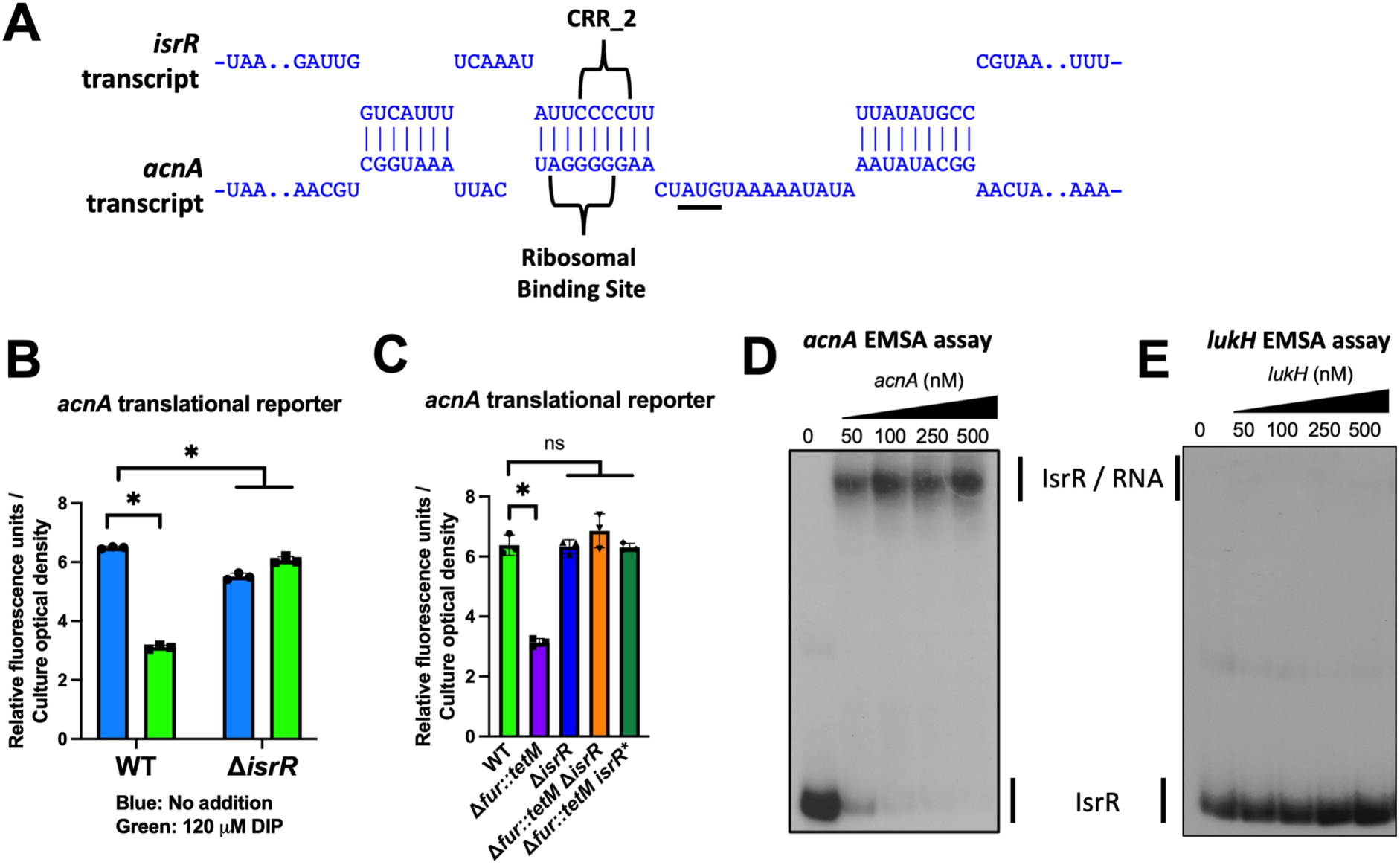
IsrR directly influences *acnA* translation. **Panel A.** IntaRNA predicted interaction between IsrR and *acnA* mRNA. Predicted interaction includes the *acnA* Shine Dalgarno and the second cytosine-rich region (CRR_2) of IsrR. The *acnA* mRNA AUG start codon is underlined. **Panel B.** Relative fluorescence of the wild type (WT) (JMB1100) and Δ*isrR* (JMB11292) strains containing the pOS_p*lgt_acnA_gfp* translational reporter after culture in TSB-Cm media with or without 120 µM DIP. **Panel C.** Relative fluorescence of the WT, Δ*isrR,* Δ*fur::tetM* (JMB10842), Δ*fur::tetM* Δ*isrR* (JMB11293), and Δ*fur::tetM isrR\** (JMB10495) strains containing the pOS_p*lgt_acnA_gfp* translational reporter after culture in TSB-Cm medium. **Panel D.** Electrophoretic mobility shift assay (EMSA) using 20,000 cpm of radiolabeled IsrR and 0-500 µM of the *acnA* transcript. **Panel E.** EMSA using 20,000 cpm of radiolabeled IsrR and 0-500 µM of the *lukH* transcript. The data displayed in panels B and C represent the average of biological triplicates with standard deviations shown. Student’s two-tailed t-tests were performed on the data and * represents a p-value of <0.05. Pictures of representative EMSA assays (n=2) are displayed in panels D and E.

To evaluate the effect of IsrR on *acnA* translation, we created a translational reporter where the constitutive *lgt* promoter drove transcription of a chimeric *acnA_gfp* generated by fusing the *acnA* 5’UTR and first two *acnA* codons in frame with the coding sequence of *gfp* (pOS_*lgt_acnA_gfp*). We used this construct to quantify *gfp* expression in the WT and Δ*isrR* strains after culture with and without DIP. The expression of *gfp* was decreased in the WT strain upon co-culture with DIP, but not in the Δ*isrR* strain (Fig 4B).

We next used the translational reporter to compare *gfp* expression in the WT, Δ*fur::tetM*, Δ*fur::tetM* Δ*isrR*, and Δ*fur::tetM isrR\** strains. When compared to the WT strain, the expression of *gfp* was decreased in the Δ*fur::tetM* strain (Fig 4C). However, the presence of either the Δ*isrR* or *isrR\** alleles reversed the Δ*fur::tetM* phenotype. These data are consistent with the hypothesis that IsrR modulates expression of the *acnA_gfp* allele *in vivo*.

We tested the hypothesis that the *isrR* and *acnA* transcripts interact *in vitro*. To this end, we performed an electrophoretic mobility shift assay (EMSA) using labeled IsrR and the 5’ of *acnA* transcript (from -41 to +541). RNA-RNA gel shifts demonstrated that titrating *acnA* transcript into samples containing IsrR decreased the rate that IsrR migrated through the polyacrylamide matrix suggesting a direct interaction between IsrR and the *acnA* transcript (Fig 4D). The *K*_d_ of IsrR for the *acnA* transcript is likely less than 50 nM. IntaRNA does not predict a strong interaction between IsrR and the *lukH* transcript, which codes for a leukotoxin. IsrR failed to interact with the *lukH* transcript using the same amounts of IsrR and *lukH* transcript as used for the *acnA* transcript EMSA (Fig 4E).

### IsrR interacts with the *acnA* ribosomal binding site

To further analyze the predicted IsrR-*acnA* mRNA interaction, we modified the *acnA* translational reporter to contain base change substitutions in and around the *acnA* Shine Dalgarno sequence, which are predicted to interact with IsrR, but still allow RBS function (pOS_*lgt_acnA1_gfp*) (Fig 5A). We hypothesized that these nucleotides were involved in IsrR-*acnA* mRNA interaction, and base substitutions would affect IsrR binding, thus decoupling *acnA_gfp* expression from IsrR control. We quantified *gfp* expression from the *acnA_gfp* and the mutated *acnA1_gfp* translational reporters in the WT, Δ*fur::tetM,* Δ*isrR,* and Δ*fur::tetM* Δ*isrR* strains. The *acnA_gfp* behaved as previously demonstrated and the Δ*fur::tetM* mutant had decreased *gfp* expression which was restored upon deletion of *isrR* (Fig 5B). The fluorescence from *acnA1_gfp* showed no significant differences between the strains examined, suggesting that the nucleotide substitutions in the *acnA* RBS decreased Fur-mediated expression control.

**Figure 5.**
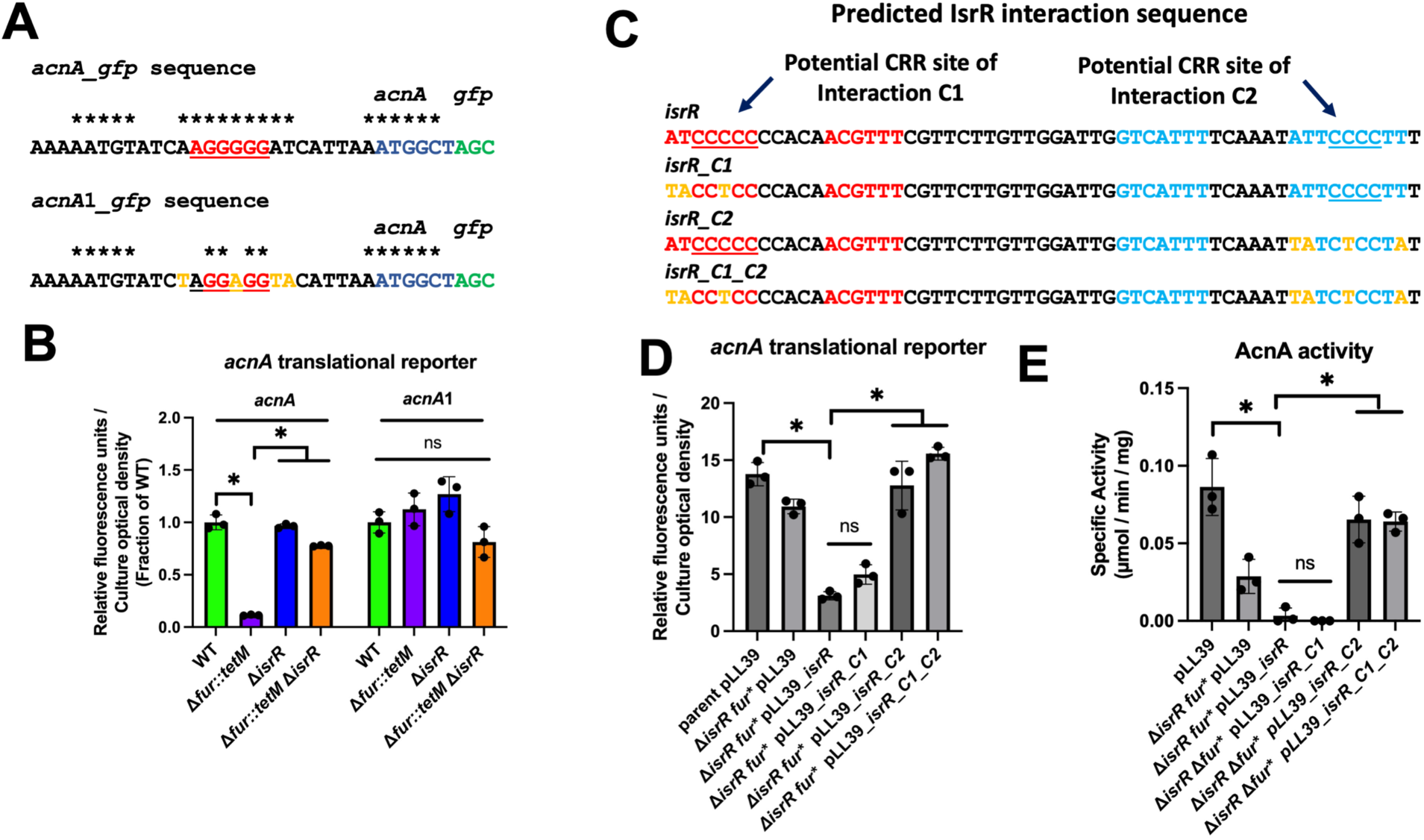
Interactions between nucleotides of the *acnA* Shine Dalgarno sequence and a cytosine rich region (CCR) of IsrR may influence IsrR-mediated *acnA* translational repression. **Panel A.** Partial sequences of the *acnA_gfp* and *acnA1*_*gfp* translational reporters. Black indicates *acnA* 5’ untranslated region (UTR), red indicates the *acnA* Shine Dalgarno sequence, blue indicates the first two codons of *acnA*, green indicates the start of the *gfp* sequence, and an asterisk above the nucleotide denotes that they are predicted to interact with IsrR (Figure 4A). Yellow nucleotides in the lower sequence indicate nucleotide substitutions in the *acnA1_gfp* translational reporter. **Panel B.** Relative fluorescence of the wild type (WT) (JMB1100), Δ*fur::tetM* (JMB10842), Δ*isrR* (JMB11292), and Δ*fur::tetM* Δ*isrR* (JMB11293) strains containing the *acnA_gfp* or *acnA1_gfp* translational reporters cultured in TSB-Cm medium. **Panel C.** Portions of IsrR and IsrR variant sequences (*isrR_C1, isrR_C2, isrR_C1_C2*) that are predicted to interact with *acnA* mRNA. Red indicates nucleotides involved in the predicted interaction that includes cytosine-rich region one (CRR_1), blue are the nucleotides in the predicted interaction with CRR_2. Underlined nucleotides indicate the IsrR C-rich regions. Yellow indicates the nucleotide substitutions on the *isrR* variants. **Panel D.** Relative fluorescence of the *1456::Tn* pLL39 (JMB11448), *fur\** Δ*isrR 1456::Tn* pLL39 (JMB11392), *fur\** Δ*isrR 1456::Tn* pLL39_*isrR* (JMB11393)*, fur\** Δ*isrR 1456::Tn* pLL39_*isrR_C1* (JMB13983)*, fur\** Δ*isrR 1456::Tn* pLL39_*isrR_C2* (JMB13984), and *fur\** Δ*isrR 1456::Tn* pLL39_*isrR_C1_C2* (JMB14314) containing the pOS_p*lgt_acnA_gfp* translational reporter. **Panel E.** Aconitase activity in cell free lysates harvested from the strains in panel D after culture in TSB medium. The data displayed in panels B, D, and E represent the average of biological triplicates with standard deviations shown. Student’s two-tailed t-tests were performed on the data and * represents a p-value of <0.05.

Coronel-Tellez et al. determined that IsrR has cytosine rich regions (CRRs) which may be used to bind mRNA targets (25). IntaRNA predicted individual interactions between the IsrR CRR_1 (Fig S5) and CRR_2 (Fig 4A) with the *acnA* transcript. We set out to determine if CRR_1 or CRR_2 is primarily responsible for driving the interactions between IsrR and the *acnA* 5’ UTR. We designed three different IsrR variants: one with mutations near CRR_1 (*isrR_C1*), one with mutations near the CRR_2 (*isrR_C2*), and one that combined both the CRR_1 and CRR_2 mutations (*isrR_C1_C2)*. The mutated *isrR* variants, as well as the wild type *isrR* were individually cloned into the pLL39 episome and integrated onto the chromosome of the Δ*isrR fur\** strain (null *fur* allele). We then used the *acnA_gfp* translational reporter to quantify the effect of the *isrR* alleles on *gfp* expression. The Δ*isrR fur\** strains carrying *isrR* or *isrR_C1* decreased *gfp* expression suggesting that both alleles function to modulate expression of *acnA_gfp* (Fig 5D). The expression of *acnA_gfp* in the Δ*isrR fur\** strains carrying the *isrR_C2* or *isrR_C1_C2* alleles phenocopied the strain carrying the empty vector suggesting that these *isrR* variants lost the ability to control *acnA_gfp* expression.

We next monitored AcnA activity in cell lysates of the WT and Δ*isrR fur\** strains carrying the empty vector or the different *isrR* alleles. As noted with the translational reporter, the Δ*isrR fur\** strains carrying *isrR* or *isrR_C1* decreased AcnA activity suggesting that both alleles function to modulate *acnA* expression (Fig 5E). The activity of AcnA in the Δ*isrR fur\** strains carrying the *isrR_C2* or *isrR_C1_C2* alleles behaved like the Δ*isrR fur\** strain carrying the empty vector suggesting that these *isrR* variants lost the ability to control *acnA* expression. Taken together these findings are consistent with the hypothesis that the CRR_2 region of IsrR is necessary to interact with the *acnA* transcript and instigate transcriptional repression.

### IsrR interacts and represses TCA cycle mRNAs

We sought to determine whether IsrR also controls the expression of genes coding for additional TCA cycle enzymes. We used IntaRNA and identified predicted IsrR interaction sites in the 5’UTR of *sdhC* (succinate dehydrogenase), *citM* (citrate synthase), and *citZ* (citrate importer) (Figs S6, S7, S8). We also identified a site in *mqo* (malate quinone oxidoreductase) in the 5’ region of the coding sequence (Fig S9). We performed RNA-RNA gel shifts to examine if IsrR could interact with the *sdh*, *mqo*, *citM*, and *citZ* transcripts *in vitro* (Figs 6A-C). Gel shifts show IsrR directly interacts with all four transcripts with an estimated *K*_d_ that is less than 50 nM.

**Figure 6.**
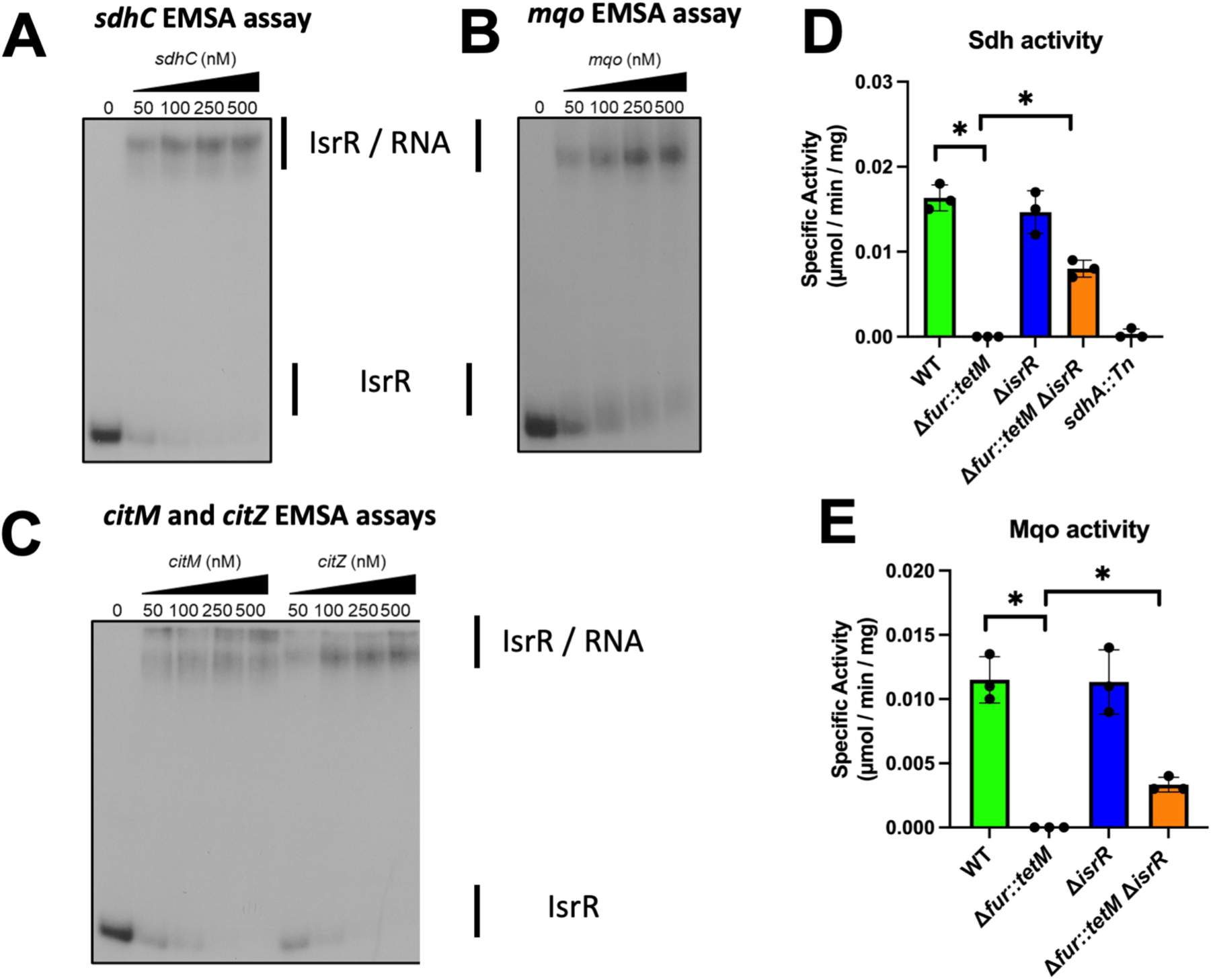
IsrR interacts with TCA cycle mRNAs and affects expression. **Panel A.** Electrophoretic mobility shift assay (EMSA) using 20,000 cpm of radiolabeled IsrR and 0-500 nM of the *sdhC* transcript. **Panel B.** EMSA using 20,000 cpm of radiolabeled IsrR and 0-500 nM of the *mqo* transcript. **Panel C.** EMSA using 20,000 cpm of radiolabeled IsrR and 0-500 nM of the *citZ* or *citM* transcripts. **Panel D.** Activity of succinate dehydrogenase (Sdh) was quantified in cell free lysates generated from the wild type (WT) (JMB1100), Δ*fur::tetM* (JMB10842), Δ*isrR* (JMB11292), and Δ*fur::tetM* Δ*isrR* (JMB11293) strains after culture in TSB medium. **Panel E.** Activity of malate quinone oxidoreductase (Mqo) in cell free lysates generated from the WT, Δ*fur::tetM,* Δ*isrR,* and Δ*fur::tetM* Δ*isrR* strains. Pictures of representative EMSA assays (n=2) are displayed in panels A-C. The data displayed in panels D and E represent the average of biological triplicates with standard deviations shown. Student’s two-tailed t-tests were performed on the data and * represents a p-value of <0.05.

We next tested the hypothesis that IsrR would negatively regulate the expression of *sdh* and *mqo*. We cultured the WT, Δ*fur::tetM,* Δ*isrR,* and Δ*fur::tetM* Δ*isrR* strains in TSB and assessed the activities of succinate dehydrogenase (Sdh) and malate quinone oxidoreductase (Mqo) in cell free lysates (Figs 6D-E). The activities of both the Sdh and Mqo were decreased in the Δ*fur::tetM* mutant and activity was partially restored upon the deletion of Δ*isrR*. Taken together, these results suggests that IsrR mediates the repression of both Fe-S cluster and non-Fe using TCA cycle enzymes in response to Fe limitation.

### IsrR impacts iron ion homeostasis

IsrR was demonstrated to downregulate expression of non-essential Fe utilizing genes including *gltA, fdh, miaB* and anaerobic nitrate respiration (*nasD* and *narG*) (25,48). This work also showed that IsrR is required for growth upon Fe depletion leading to a model wherein IsrR contributes to Fe sparing upon Fur demetallation. TCA cycle enzymes are a cellular Fe sink, and herein, we demonstrate that IsrR represses the expression of TCA cycle genes in the absence of Fur, which contributes to the proposed Fe sparing model.

We tested the hypothesis that *isrR* expression promotes Fe uptake and/or an increase in Fe ions not ligated by macromolecules (also called “free Fe”). The antibiotic streptonigrin when combined with intracellular Fe(II) and a reducing agent promotes killing by causing double stranded DNA breaks (3,7). Therefore, increased killing by streptonigrin is correlated with increased pool of Fe that is not chelated by macromolecules. We assayed streptonigrin sensitivity by spotting streptonigrin on top agar overlays containing the WT, Δ*fur::tetM,* Δ*isrR,* or Δ*fur::tetM* Δ*isrR* strains. The Δ*fur::tetM* strain displayed increased streptonigrin sensitivity compared to the WT (Fig 7A). This phenotype was expected since a Δ*fur* mutant has derepressed Fe uptake (42). The Δ*fur::tetM* Δ*isrR* had decreased streptonigrin sensitivity when compared to Δ*fur::tetM,* suggesting a role for IsrR in increasing streptonigrin sensitivity on a Δ*fur::tetM* strain. We confirmed a role for IsrR in increasing streptonigrin sensitivity by inducing the expression of *isrR* or an antisense *isrR* (as_*isrR*) in the WT strain using the pEPSA5 vector. Induced expression of *isrR*, but not as_*isrR*, resulted in increased streptonigrin sensitivity. These data are consistent with the hypothesis that upon Fe limitation and Fur derepression, *isrR* expression increases the pool of free intracellular Fe.

**Figure 7.**
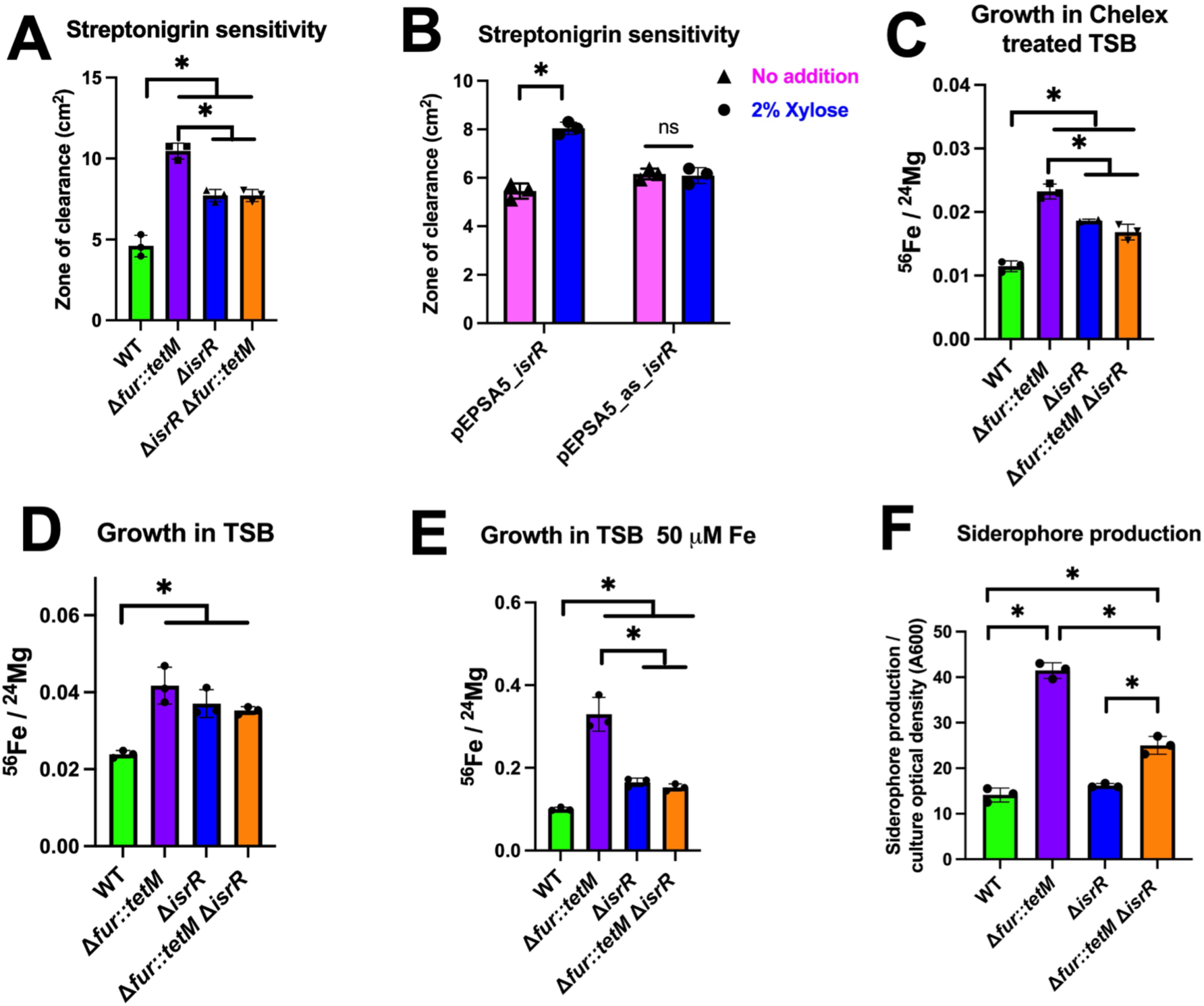
IsrR impacts iron homeostasis. **Panel A.** The wild type (WT)(JMB1100), Δ*fur::tetM* (JMB10842), Δ*isrR* (JMB11292), and Δ*fur::tetM* Δ*isrR* (JMB11293) strains were plated as top agar overlays on solid TSA, followed by spotting 2.5 µg streptonigrin. **Panel B.** The WT strain containing pEPSA5_*isrR* or pEPSA_as_*isrR* were plated as top agar overlays on solid TSA-Cm with or without 2% xylose, followed by spotting 2.5 μg streptonigrin. For Panels A and B, the zones of clearance resulting from streptonigrin growth inhibition was quantified. **Panels C, D, and E.** The ratio of ^56^Fe and ^24^Mg abundances were quantified in whole cells using ICP-MS after culture in Chelex treated TSB (Panel C), TSB (Panel D), or TSB supplemented with 50 µM Fe (Panel E). The ratio of ^56^Fe / ^24^Mg is displayed for WT, Δ*fur::tetM,* Δ*isrR,* and Δ*fur::tetM* Δ*isrR* strains. **Panel F**. Siderophore production from spent culture supernatants from the WT, Δ*fur::tetM,* Δ*isrR,* and Δ*fur::tetM* Δ*isrR* strains was quantified. The data displayed represent the average of biological triplicates with standard deviations shown. Student’s two-tailed t-tests were performed on the data and * represents a p-value of <0.05.

We next evaluated the role of IsrR in Fe ion uptake. We quantified total ^56^Fe pools using Inductively Coupled Mass Spectrometry (ICP-MS) after growth in 1) TSB, 2) TSB with 50 µM Fe(II), and 3) TSB treated with Chelex to decrease the titers of divalent metals (Figs 7C, 7D, 7E). The Δ*fur::tetM*, Δ*isrR*, and Δ*fur::tetM* strains had increased titers of ^56^Fe compared to the WT in all three media. However, the Δ*isrR* and Δ*fur::tetM* Δ*isrR* strains had decreased ^56^Fe levels compared to Δ*fur* in Chelex-treated TSB and TSB supplemented with Fe. The intermediate iron levels of the Δ*fur::tetM* Δ*isrR* suggest that IsrR contributes to the increased Fe levels in strains lacking Fur.

Lastly, we examined whether IsrR has a role in siderophore production. We quantified total siderophore production in the WT, Δ*fur::tetM,* Δ*isrR,* and Δ*fur::tetM* Δ*isrR* strains after culture in Chelex-treated TSB. As expected, a Δ*fur::tetM* had increased siderophore production compared to the WT (Fig 7F). The Δ*fur::tetM* Δ*isrR* double mutant strain produced fewer siderophores than the Δ*fur::tetM*, but more than the Δ*isrR* and WT strains. Taken together, these findings verify a role for IsrR in the Fe ion homeostasis and suggest that the role is, in part, the result of decreased siderophore production.

### IsrR and Fur contributes to *S. aureus* pathogenesis

To define the respective roles of IsrR and Fur in pathogenesis we first tested their roles in a model of acute pneumonia. In both bronchoalveolar lavage fluid (BALF) and lung tissue we observed the importance of IsrR on pathogenesis (Fig. 8A&B). While we did not observe a significant impact of Fur in this model, in either the WT or *fur* background, inactivation of *isrR* led to decrease in bacterial survival. In BALF, inactivation of *isrR* led to a nearly 40-fold reduction in bacteria, this held true in the lung with a 25-fold decrease. We next sought to determine how the absence of IsrR or Fur would impact skin infection. To this end, we performed a murine model of skin infection and monitored several outcomes. We observed no difference in lesion size between the Δ*isrR* mutant and WT strain. By contrast, there was a significant decrease in lesion size for the Δ*fur::tetM* and Δ*fur::tetM* Δ*isrR* mutants compared to the WT-infected mice. While lesion size reports the overall surface lesion, necrosis size quantifies fully necrotic tissue that forms during infection and is observed as a scab-like structure. The same trends were observed for necrosis between the strains.

**Figure 8.**
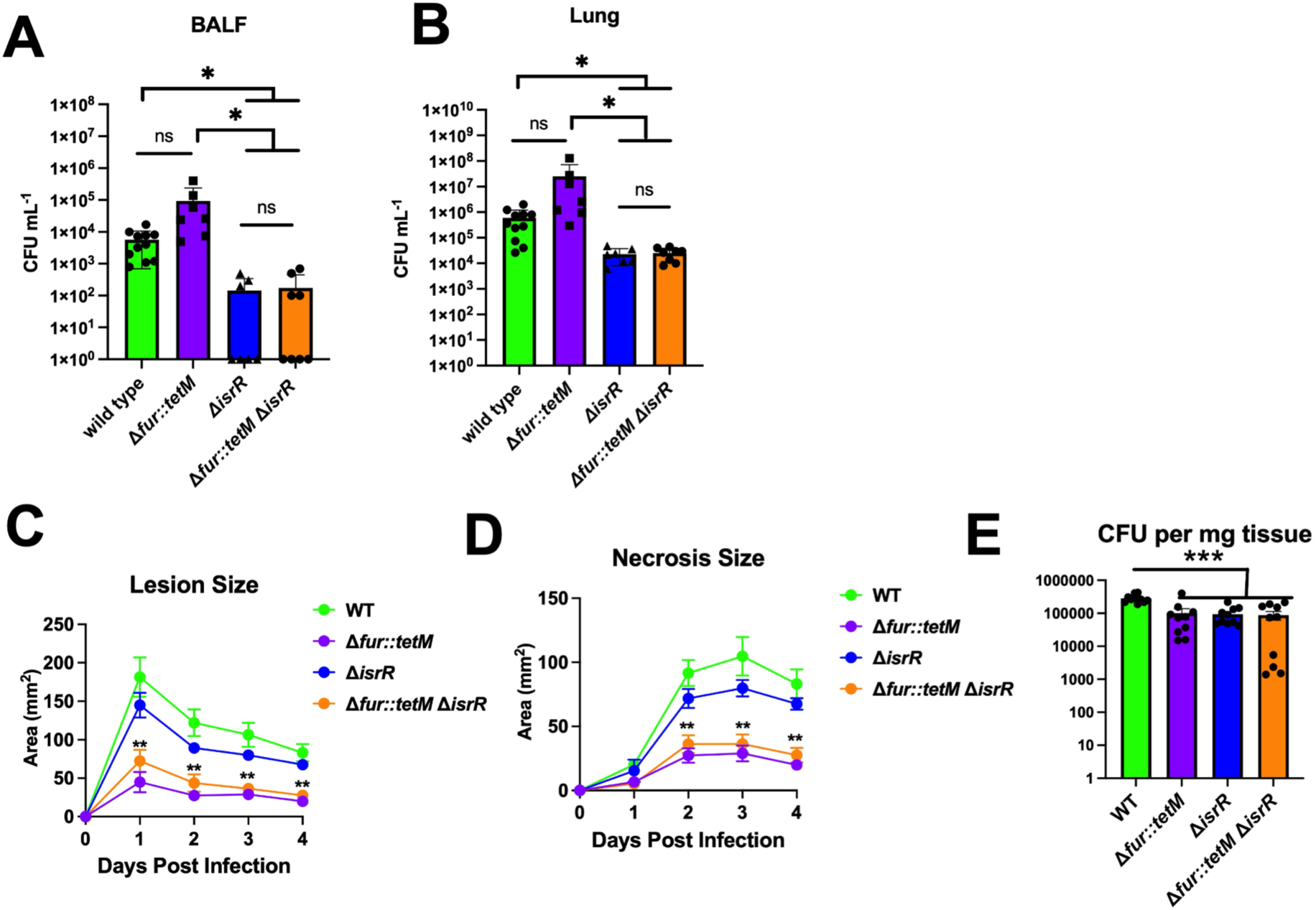
IsrR and Fur are important for tissue damage and colonization during infection. **Panels A** and **B.** The wild type (WT)(JMB1100), Δ*fur::tetM* (JMB10842), Δ*isrR* (JMB11292), and Δ*fur::tetM* Δ*isrR* (JMB11293) strains were tested in a model of acute pneumonia infection. Data represent bacterial counts 24 hours after intranasal infection in bronchoalveolar lavage fluid (Panel A) and lung tissue (Panel B). **Panels C and D**, the wild type (WT), Δ*fur::tetM,* Δ*isrR,* and Δ*fur::tetM* Δ*isrR* strains were injected subcutaneously into C57BL/6J mice and total lesion size (Panel C) and necrosis size (Panel D) were monitored over time. **Panel E**. Local bacterial titers were quantified four days post infection. Data is a representative experiment with n=10. For A and B, each symbol represents the mean. For C and D, each dot is an individual animal, and the bar or line represents the mean. Error bars represent the SEM and may be smaller than symbols. * and ** indicates p<0.05 and p<0.01, respectively, compared to WT for Δ*fur::tetM* and Δ*fur::tetM* Δ*isrR* by Mann-Whitney test.

**Figure 9.**
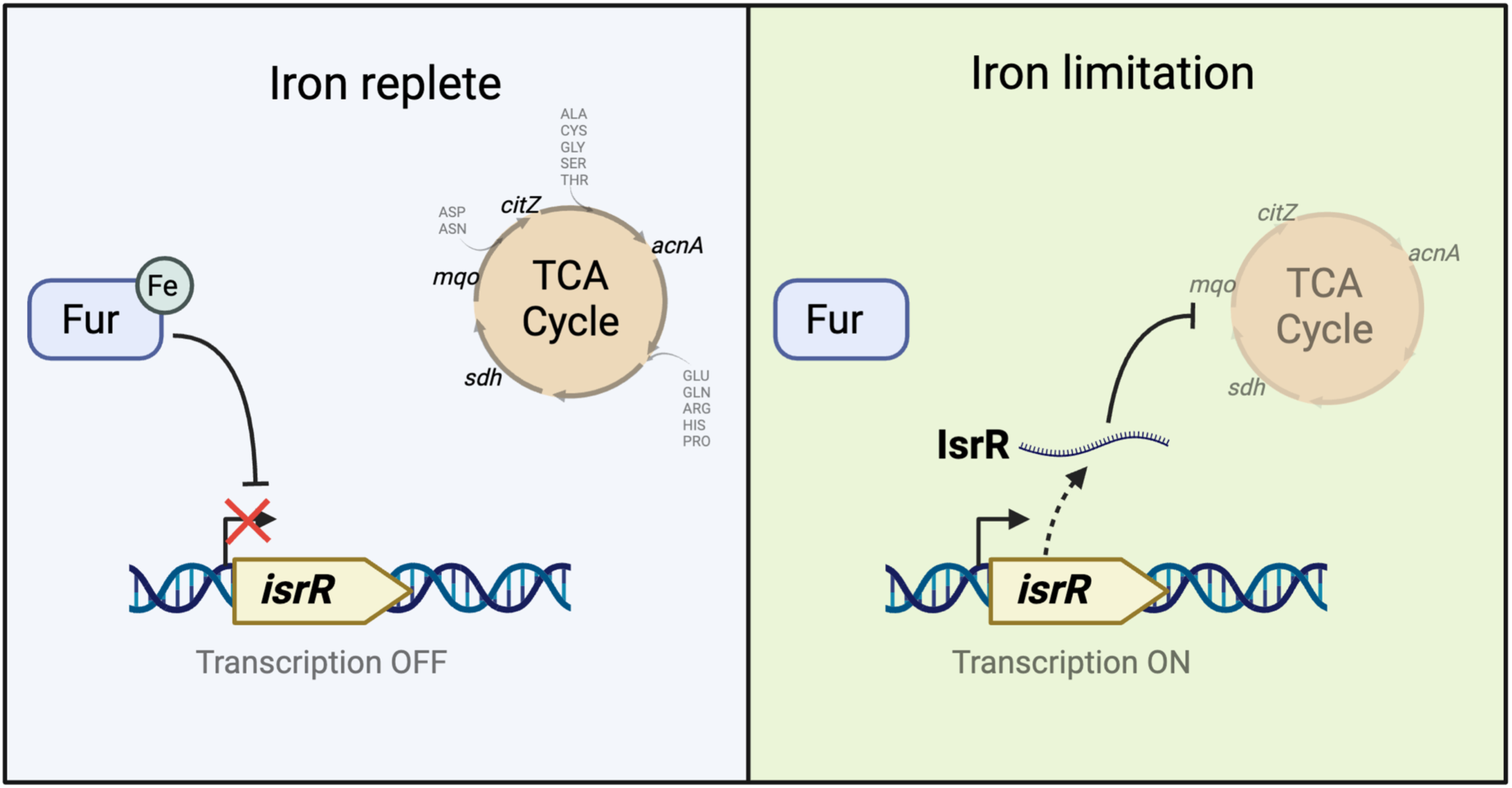
Model for IsrR-dependent regulation of TCA cycle expression. Growth in iron ion replete conditions results in metalation of Fur and transcriptional repression of *isrR*, which produces a non-coding small regulatory RNA. Upon iron limitation, Fur is demetallated, and *isrR* is expressed. IsrR forms complexes with the *acnA*, *sdh*, *citM*, and *citZ* mRNA transcripts resulting in decreased expression and decreased carbon flux through the TCA cycle. Decreased expression of TCA cycle enzymes results in an inability to grow using amino acids for carbon and energy. Figure was created using Biorender.

To determine if reduced lesion or necrosis size in the absence of Fur was due to changes in bacterial colonization, bacterial titers at the site of infection were determined at 4 days post-infection. We observed an ∼3-fold decrease in bacterial titers in all mutant strain-infected mice compared to WT-infected mice at this time point. Since the Δ*isrR* mutant showed reduced bacterial numbers but not decreased lesion formation, we interpret this to mean that the smaller lesions observed in the Δ*fur::tetM* and Δ*fur::tetM* Δ*isrR* mutants are not likely due to changes in bacterial titers at the site of infection.

Tissue damage likely results from a combination of bacterial factors and the host immune response. Considering the reduced tissue damage in the absence of Fur, we measured cytokine and chemokine levels at the site of infection to provide insight into the immunological changes occurring during infection with *fur* mutants. We used a panel of proinflammatory cytokines that are important for immune cell recruitment and/or activation. We did not observe a change in KC, MCP-1, GM-CSF, or MIP1α between any of the strains. Thus, tissue damage differences did not correlate with any changes in these chemokines and cytokines. IL-1β, G-CSF, and IL-6 were decreased in all mutant bacterial infections. In contrast, we observed increased IL-1α but decreased TNFα and G-CSF in a Fur-dependent manner and these changes correlate with decreased lesion size when Fur was absent. We did not perform statistical analysis on CCL5 (RANTES), but while it was readily detectable in wild type-infected mice, most mutant mice had levels below the limit of quantification. The finding that some cytokine levels, but not others, differed when Fur was absent suggests specific immunological changes are occurring during infection in a Δ*fur::tetM* mutant. What those changes are in immune cell recruitment or activation and the mechanism by which this occurs will require additional investigation.

## Discussion

This study was initiated to understand why a strain lacking Fur has decreased TCA cycle function. Work by others led to the hypothesis that growth in Fe limiting conditions decreases the levels of Fe-bound Fur, resulting in altered affinity for DNA, and derepression of the Fur regulon (11,18). Bioinformation work led us to predict that the genes that Fur directly regulates are utilized for Fe uptake, as well as a gene for Fe storage (*dps*) (42). However, we discovered that a Δ*fur* mutant had greatly reduced AcnA activity. We also noted that neither Δ*fur* or *acnA::Tn* mutants could grow using amino acids as carbon or energy sources providing us with a phenotype that we could exploit.

We used a suppressor screen approach to identify IsrR as the Fur-regulated negative regulator of aconitase expression. To our knowledge, this is the first case of a suppressor screen identifying a sRNA through the suppressive effects of a null mutation. The suppressor mutations in *isrR* (called *isrR\**) allowed for increased growth of the Δ*fur* mutant on defined amino acid media. Interestingly, all fifteen of the *isrR\** mutant strains, isolated during two separate screening events, contained the same mutation in the operator of *isrR* suggesting that there is something unique about this allele. In alternate organisms, small proteins, such as Hfq in *E. coli*, promote interaction between a sRNA and a target mRNA (49). The finding that suppressor mutations only mapped to the promoter of *isrR*, and not to alternate loci including *hfq*, suggest that *S. aureus* does not require a single protein to facilitate interaction between IsrR and target RNAs. The study by Coronel-Tellez *et al*. also found that a strain with a deletion mutation in the gene predicted to code the Hfq homologue did not alter IsrR-dependent regulation of expression (25). Alternatively, there could be more than one chaperone that aids IsrR regulation that share functional overlap.

The *isrR* operator has two Fur-boxes and the *isrR\** mutation is in the Fur-box that is proximate to the transcription start site (25). The mutation resulted in a loss of transcription and possibly Fur-mediated genetic regulation. It also resulted in an inability for *isrR* transcription to be induced during low Fe. It is currently unknown how this mutation decreased transcription, but it did change the sequence of a proposed -35 sigma factor A recognition sequence from TTGAAA to TTTAAA (consensus is: TTGATA). The *isrR\** mutation did not alter *isrR* transcript degradation, and strains expressing *isrR\** or *isrR* using a non-native promoter resulted in phenotypic similarities suggesting that *isrR\** is functional *in vivo*, but the mutation decreases transcription.

The AcnA activity of a Δ*fur* mutant was increased by the deletion of *isrR* and over-expression of *isrR* in the WT decreased AcnA activity. We identified a predicted IsrR binding site in the *acnA* transcript and demonstrated that IsrR interacted with the *acnA* transcript *in vitro*. The predicted interaction site overlayed the *acnA* Shine Dalgarno sequence. Introducing mutations into the Shine Dalgarno sequence of *acnA* that preserve the ribosomal binding site, but impact predicted base-pairing with IsrR, decreased IsrR mediated control of *acnA* expression. Additionally, mutations impacting the second cytosine rich region (CCR_2) of IsrR predicted to interact with the *acnA* transcript also decreased IsrR control over *acnA* expression. These data are consistent with the model wherein IsrR binds the *acnA* transcript and mediates translational repression through occlusion of the RBS. This model is supported by previous findings where IsrR mediates translational repression of *fdhA* and *gltB2*, which have similar predicted IsrR pairing sites as the *acnA* transcript, and don’t appear to trigger mRNA degradation (25). The introduction of the Δ*isrR* mutation to the Δ*fur::tet* strain resulted in partial recovery of activity suggesting some level of Fur-mediated control of *acnA* expression. The promoter of *acnA* contains a Fur box terminating at position -179. We are currently trying to determine if Fur itself acts as an activator of acnA transcription.

Further examination found that IsrR binds to additional mRNA transcripts that code for enzymes of the TCA cycle including *sdh, mqo, citM* and *citZ*. Activity assays were used to verify that Mqo and Sdh activity was decreased in a Δ*fur* mutant, and this regulation was, in part, relieved by the deletion of *isrR*.

The results herein in combination with previous findings demonstrate that IsrR represses expression of mRNA coding the Fe-requiring proteins AcnA, Sdh, Mqo, GltB2, FdhA, MiaB (25,48). Coronel-Tellez *et al*. found that an Δ*isrR* mutant had decreased growth when challenged with DIP on solid media supporting the model wherein IsrR functions to spare Fe in *S. aureus* (25). To further support this model, we examined intracellular Fe ion pools. As previously witnessed, a Δ*fur* mutant was more susceptible to killing by streptonigrin, suggesting an increased free Fe pool (7). This sensitivity of the Δ*fur* mutant was lessened by deletion of *isrR*, but not returned to the levels seen in WT. These data are consistent with a model wherein *isrR* expression promotes an increased Fe ion pool that is not ligated by macromolecules; however, it is currently unknown if this is the result of decreased expression of Fe-requiring proteins or increased Fe uptake or both phenomena. The expression of *isrR* in the Δ*fur* mutant also increased total Fe pools when *S. aureus* was cultured in rich complex medium under Fe deplete or replete conditions. Production of *S. aureus* siderophore staphyloferrin B (Sbn) requires citrate synthesized from o-phospho-L-serine and glutamate (50). It is tempting to speculate that IsrR decreases TCA cycle function to decrease glutamate catabolism and promote citrate usage for Sbn synthesis. Consistent with this speculation, introduction of a Δ*isrR* mutation into the Δ*fur* mutant decreased siderophore production. It is currently unknown why the Δ*isrR* strain had increased free Fe and cell associated Fe compared to the WT strain. The simplest explanations are that 1) growth in the TSB media utilized resulted in some Fur-dependent derepression (low Fe titers, ROS accumulation, etc.) of *isrR*, 2) *isrR* is regulated by an alternate transcriptional regulator(s), or 3) IsrR has pleiotropic roles in promoting and inhibiting expression of different Fe uptake mechanisms. Further studies are necessary to parse out the role of IsrR on regulation of individual Fe uptake systems. So far, IsrR has been shown to regulate ten mRNAs that encompass nitrogen homeostasis (*gltB2, narG, nasD*), fermentation (*fdhA*), tRNA modification (*miaB*), and TCA cycle (*acnA, sdhC, mqo, citM, citZ*) (25,48). IsrR likely functions to repress the expression of these enzymes to spare Fe for use by alternate proteins essential for fitness; however, it is unclear what these proteins are. Fe-S cluster synthesis is essential in *S. aureus*, but the essential Fe-S protein(s) remain elusive (7).

The link between Fe, Fur, and metabolism has been observed across bacteria, including *S. aureus.* This relationship was first explained by the discovery of the Fur-regulated sRNA RyhB in *E. coli* (*23,51*). RyhB expression results in an Fe sparing response where non-essential Fe-using proteins involved in processes like the TCA cycle (*acnB, sdh, fumA*), iron storage (*ftnA, bfr*), oxidative stress (*sodB*), respiration (*nuo*), and Fe-S cluster assembly (*isc*) are downregulated in response to Fe limitation (23). RyhB homologs have been found in other enterobacteria such as *Salmonella, Shigella, and Yersinia,* and functional analogs have been described in *Pseudomonas* (PrrF1 and PrrF2) and *B. subtilis* (FsrA) (22,52–55). As Coronel-Tellez *et al*. indicated, it is remarkable that IsrR, although not similar in sequence to these alternate sRNA, functions in a homologous manner to decrease expression of Fe requiring enzymes and processes (25). Moreover, expression of these alternate sRNA is controlled by Fur.

The virulence defects of the *isrR* mutants may suggest a proper Fe sensing and response is required for full host colonization. The importance of Fe homeostasis is highlighted by results showing the colonization defects of strains lacking Fur or IsrR in lung and skin murine models of infection. IsrR was also shown to be required for full lethality of *S. aureus* in a mouse septicemia model (5). A *fur* mutant in the Newman genetic background had increased exoprotein protein, leukotoxin production, hemolysis production, and increased killing of HL-60 cells. However, the fur mutant also had decreased survival in a neutrophil killing assay and in a model of murine pneumonia. Interestingly, depletion of neutrophils nullified the defective lung colonization of the *fur* mutant suggesting that the increased susceptibility of the fur mutant to neutrophil killing was contributing to its inability to colonize lung tissue (5). This contrasts with our results that did not observe a decrease in pathogenesis with the *fur* mutant. This might be due to the different strains of *S. aureus* used (USA300_LAC and Newman), as well as the earlier study relying on a higher infection dose.

The effect of IsrR on virulence could be indirect and a consequence of metabolite imbalance. Altering metabolic status can impact metabolite pools, which can be sensed by transcriptional regulators that control virulence factor production (56,57). For example, TCA cycle repression could alter GTP and branched-chain amino acid pools (BCAAs) which are sensed by CodY resulting in altered virulence factor production (58,59). Likewise, IsrR-mediated TCA cycle repression could increase pyruvate titers. Increased pyruvate upregulates virulence factor production in *S. aureus* through complex regulatory networks involving the Agr, Arl, and Sae regulatory systems (60). Further study into the regulatory effects of IsrR should reveal how IsrR contributes to *S. aureus* virulence and will shed light on the cellular response of *S. aureus* to host encountered Fe limitation.

Thanks to high throughput sequencing technologies, hundreds of sRNA candidates have been identified in *S aureus*. Nevertheless, only 50 sRNAs fit the requirements to be *bona fide* trans acting sRNAs and around fifteen have been associated with their mRNA targets and biological functions (35). The studies in this manuscript have furthered our understanding of one of these *bona fide* sRNA. The results presented demonstrate that expression of *isrR* modulates the expression of the TCA cycle and directly interacts with mRNA transcripts. We also demonstrate that both IsrR and Fur contribute to cellular Fe homeostasis and virulence. Future studies will investigate the mechanisms by which IsrR and Fur alter pathogenesis.

## Data Availability Statement

The data underlying this article are available in this article and in its online supplementary material. Strains and plasmids will be made available upon request.

