## Supplemental Data for "The *Staphylococcus aureus* small non-coding RNA IsrR regulates TCA cycle activity and virulence"

Table S1. DNA primers used in this study

| Name | Sequence |
|---|---|
| Tsr25 qPCR fwd | ACGTTTCGTTCTTGTTGGATTG |
| Tsr25 qPCR rev | GTGTCGTAAGGGTTTACTGCT |
| 16S qPCR set3 | TGAGTGCAGAAGAGGAAAGTG |
| 16S qPCR set3 | CGTCAGTTACAGACCAGAAAGT |
| Tsr25 comp 5 SalI | CCCGTCGACGTGTCAGATGACATTAAATAGCATCTCC |
| Tsr25 comp 3 BamHI | CCCGGATCCCCGCAATATTAATTGTTGTCATATAC |
| Tsr25 pEPSA 5 EcoRI | GGGGAATTCTAATAGTAGTTGAAAATGATTATC |
| Tsr25 pEPSA 3 SalI | GGGGTCGACTAAAAGCAGTAAACCTAAAGTGTCG |
| Tsr25* pEPSA 5 EcoRI | GGGGAATTCTAATAGTAGTTTAAAATGATTATC |
| Tsr25 trunk pEPSA 5 EcoRI | GGGGAATTCAAAATGATTATCAATACCATA |
| tsr25 pMAL 5 SalI | GGGGTCGACTAATAGTAGTTGAAAATGATTATC |
| tsr25 pMAL 3 EcoRI | GGGGAATTCTAAAAGCAGTAAACCTAAAGTGTCG |
| tsr25 AF | CAAGAGCTCCGTTCTTATCAATTGTAATACC |
| tsr25 AR | CGGACGCGTCATTAGTGAGAATCATTGTC |
| tsr25 BF | CGGACGCGTGCTTTTGTTTTTTGTAAAAAGAAAACC |
| tsr25 BR | GGGGTACCCGTTTAATTGATATAACAGG |
| pOStsr25_2forHindIII | GGGAAGCTTGCGTAATCATGCGTTTTATT |
| pOStsr25_2revkpnI | GCGGGTACCCCAACAAGAACGAAACGTTGTG |
| Tsr25del up For | CAAGAGCTCCGTTCTTATCAATTGTAATACC |
| Tsr25del up Rev | CGGACGCGTCATTAGTGAGAATCATTGTC |
| Tsr25del dwn For | CGGACGCGTGCTTTTGTTTTTTGTAAAAAGAAAACC |
| Tsr25del dwn Rev | GGGGTACCCGTTTAATTGATATAACAGG |
| pEPSA5_acnAEcoRI | CCCGAATTCATCAAGGGGGATCATTAAATGGCTGCAAATTTTAAAGAGCAATC |
| pEPSA_acnA3SalI | CCCGTCGACGAGCACTATCAAAGTGCC |
| Plgt_acnA | GAGTAGGGATAAATACAATTGAGGTGAACATATACTCTGATTAAAAAGTCAAAAC |
| GfP_plgt rev (gfp amplicon) | AAAGGGGGAAACACTACCCCCTTGTTTGGATCTTAGTGGTGGTGGTGGTGGTGGG AT |
| acnA_gfp Wt A rev (acnA amplicon) | GTGAAAAGTTCTTCTCCTTTGCTAGCCATTTAATGATCCCCCTTGATACATTTT |
| acnA_gfp WtA for (gfp amplicon) | AAAATGTATCAAGGGGGATCATTAAATGGCTAGCAAAGGAGAAGAACTTTTCAC |
| acnA_gfp Mut C Rev (acnA amplicon) | GTGAAAAGTTCTTCTCCTTTGCTAGCCATTTAATGTACCTCCTAGATACATTTTTA |
| acnA_gfp Mut C For (gfp amplicon) | TAAAAATGTATCTAGGAGGTACATTAAATGGCTAGCAAAGGAGAAGAACTTTTCAC |
| isrR5north | GTTCTTGTTGGATTGGTCAT |
| isrR3north | GTGTCGTAAGGGTTTACTGC |
| T7-*isrR*-For | TAATACGACTCACTATAGGGAAAATGATTATCAATACCAC |
| T7-*isrR*-Rev | AAACAAAAGCAGTAAACCTAAAGTGTCG |
| T7-*acnA*-For | TAATACGACTCACTATAGGGAAGGCATATAAATATAAAAATGTATC |
| T7-*acnA*-Rev | GAACTATTCCAGTTGCAGGAGG |
| T7-*citM*-For | TAATACGACTCACTATAGGGAGGGGGATTATGTAAATTG |
| T7-*citM*-Rev | CTTACATCTTGATCTCGTTCATATAC |
| T7-*citZ*-For | TAATACGACTCACTATAGGGAAAGCCATTTCATAAAGAAAC |
| T7-*citZ*-Rev | GACCAGCGTCTTTCGTAATTTG |
| T7-*mqo*-For | TAATACGACTCACTATAGGGAGTAATCAGCAATCAAATTAAATTG |
| T7-*mqo*-Rev | GATCTGGGTTATCTAATTGTCCTG |
| T7-*sdhC*-For | TAATACGACTCACTATAGGGAACAAAGTTATTTTTTTGAATGTACG |
| T7-*sdhC*-Rev | TTACATAAAGGCAATAATTGCAGTAACAC |
| T7-*lukH*-For | TAATACGACTCACTATAGGGATTTAATAAATAGTTAAATATATATTCTCTG |
| T7-*lukH*-Rev | GCCACTTCTTACTAATGCTGGG |

Table S2. synthetic DNA used to generate the mutant *isrR* alleles.

| | |
|---|---|
| *isrR_C1* | GTTTCTAATTGACAATGATTCTCACTAATGTATAATAGTAGTTGAAAA<br>TGATTATCAATACCATATAGAACTACCTCCCCACAACGTTTCGTTCTT<br>GTTGGATTGGTCATTTTCAAATATTCCCCTTTTATATGCCCGTAAAAG<br>ACAATATACGTTATAACAACGTTTTATAAAAGCAGTAAACCCTTACGA<br>CACTTTAGGTTTACTGCTTTTGTTTTTTGTAAAAAGAAAACCATACGC<br>TATGCGTATGGTTCAGAAAA |
| *isrR_C2* | GTTTCTAATTGACAATGATTCTCACTAATGTATAATAGTAGTTGAAAA<br>TGATTATCAATACCATATAGAACATCCCCCCCACAACGTTTCGTTCTT<br>GTTGGATTGGTCATTTTCAAATTATCTCCTATTATATGCCCGTAAAAG<br>ACAATATACGTTATAACAACGTTTTATAAAAGCAGTAAACCCTTACGA<br>CACTTTAGGTTTACTGCTTTTGTTTTTTGTAAAAAGAAAACCATACGC<br>TATGCGTATGGTTCAGAAAA |
| *isrR_C1_C2* | GTTTCTAATTGACAATGATTCTCACTAATGTATAATAGTAGTTGAAAA<br>TGATTATCAATACCATATAGAACTACCTCCCCACAACGTTTCGTTCTT<br>GTTGGATTGGTCATTTTCAAATTATCTCCTATTATATGCCCGTAAAAG<br>ACAATATACGTTATAACAACGTTTTATAAAAGCAGTAAACCCTTACGA<br>CACTTTAGGTTTACTGCTTTTGTTTTTTGTAAAAAGAAAACCATACGC<br>TATGCGTATGGTTCAGAAAA |

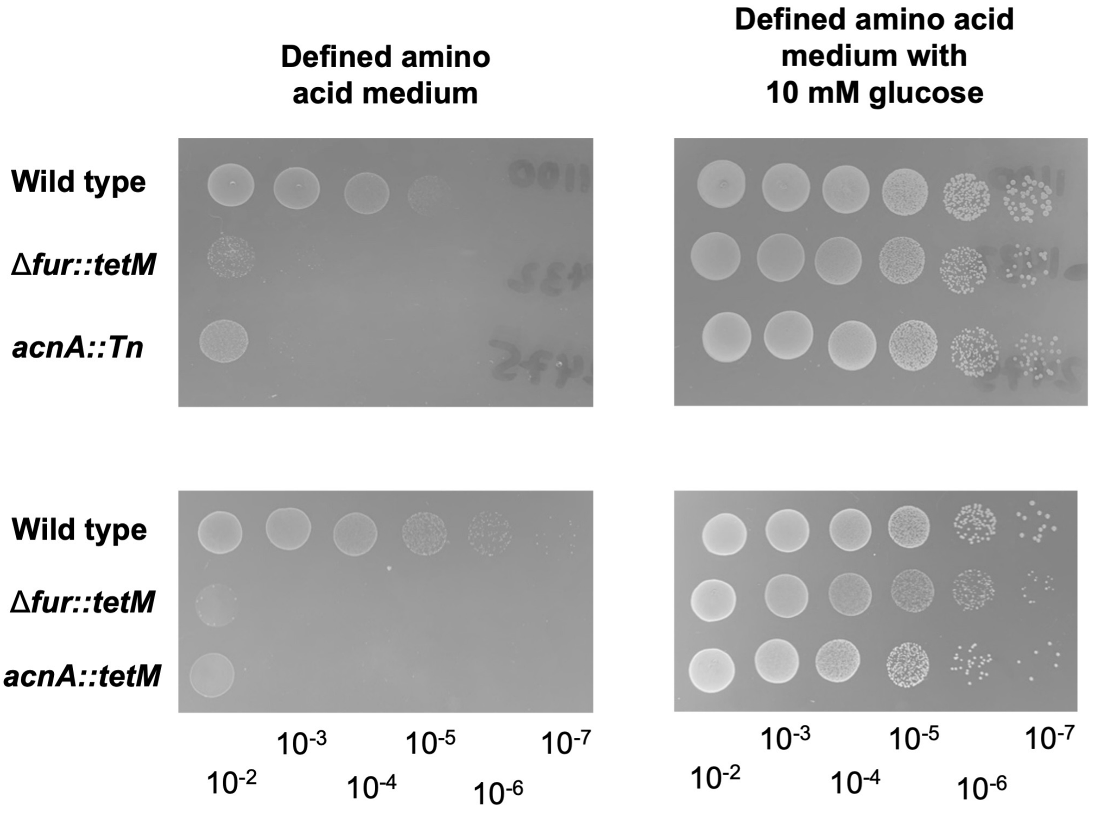


**Supplemental Figure 1.** *Staphylococcus aureus* strains lacking Fur or AcnA have a growth defect on amino acid medium. The wild type (JMB1100), Δ*fur::tetM* (JMB10842), and *acnA::Tn* (JMB11803) strains were standardized to an optical density ($A_{600}$) of 2, serial diluted (10-fold per dilution), and spotted on solid defined amino acid medium with or without 10 mM glucose. A photo of a representative experiment is shown.

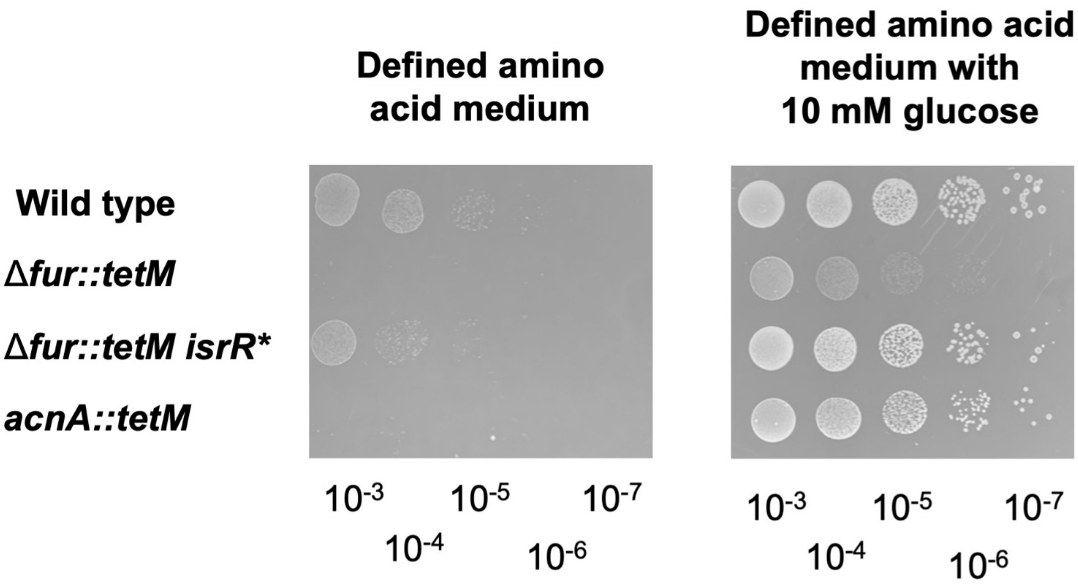


**Supplemental Figure 2.** The *isrR** mutation causes increased growth of a Δ*fur* mutant on solid chemically defined media. The wild type (JMB1100), Δ*fur::tetM* (JMB10842), Δ*fur::tetM isrR** (JMB10495), and *acnA::Tn* (JMB11803) strains were standardized to an optical density ($A_{600}$) of 2, serial diluted (10-fold per dilution), and spotted on solid defined amino acid medium with or without 10 mM glucose. A photo of a representative experiment is shown.

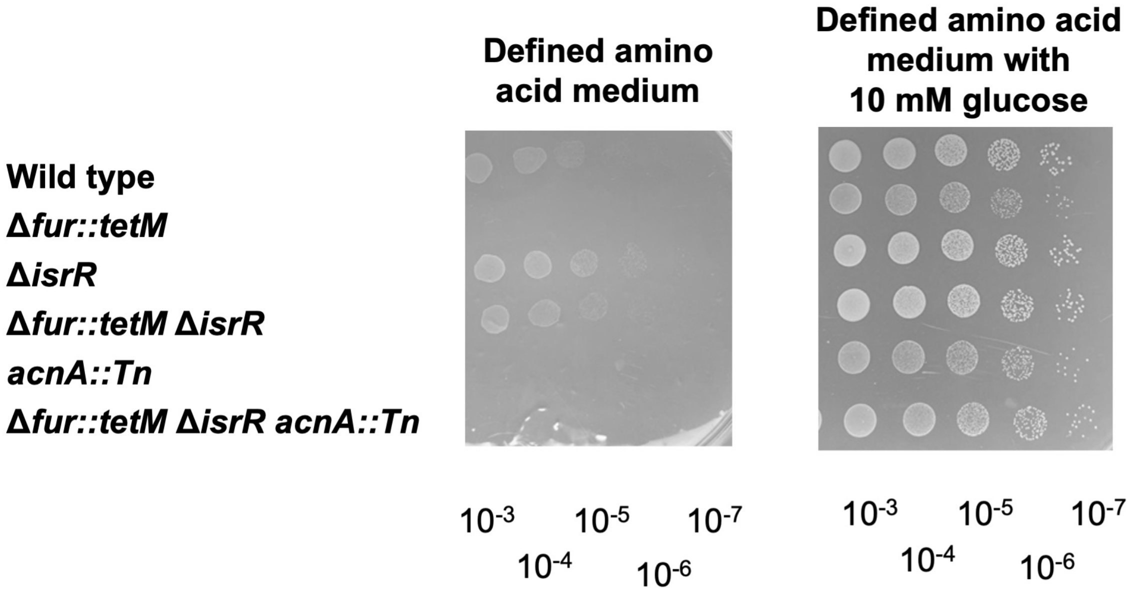


**Supplemental Figure 3.** Deletion of *isrR* leads to increased growth of a Δ*fur* mutant on chemically defined media. The wild type (JMB1100), Δ*fur::tetM* (JMB10842), Δ*isrR* (JMB11292), Δ*fur::tetM* Δ*isrR* (JMB11293), *acnA::Tn* (JMB11803), and Δ*fur::tetM* Δ*isrR* *acnA::Tn* (JMB11806) strains were standardized to an optical density($A_{600}$) of 2, serial diluted (10-fold per dilution), and spotted on solid defined amino acid medium with or without 10 mM glucose. A photo of a representative experiment is shown.

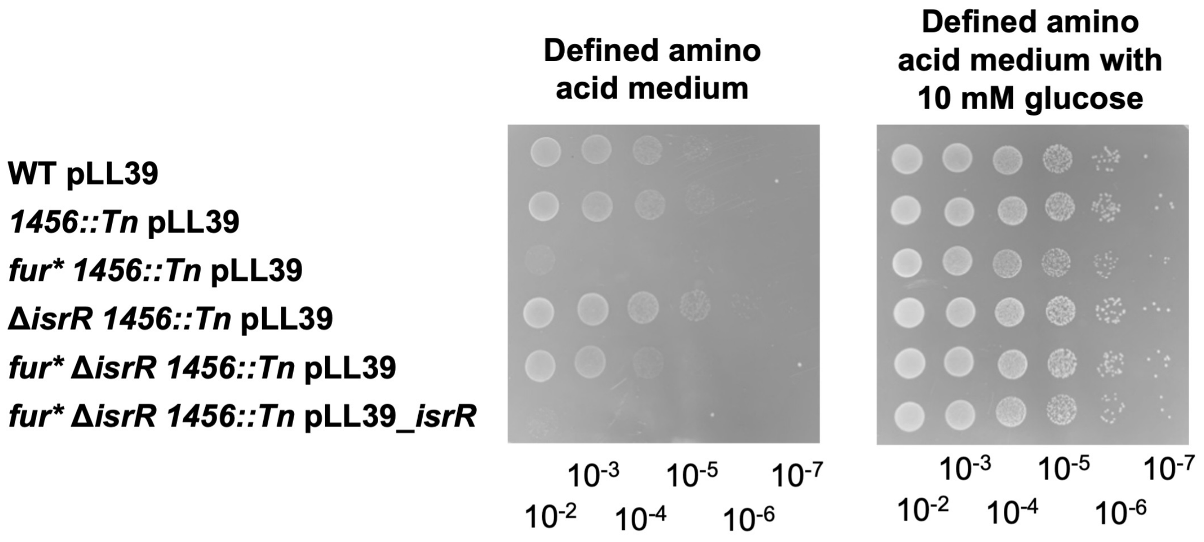


**Supplemental Figure 4.** Genetic complementation of the Δ*isrR* mutation. The wild type with pLL39 (JMB1886), *1456::Tn* pLL39 (JMB11448), *fur* 1456::Tn* pLL39 (JMB11449), Δ*isrR 1456::Tn* pLL39 (JMB11395), *fur** Δ*isrR 1456::Tn* pLL39 (JMB11392), and *fur** Δ*isrR 1456::Tn* pLL39_*isrR* (JMB11393) strains were standardized to optical density of 2 ($A_{600}$ nm), serial diluted (10-fold per dilution), and spotted on solid defined amino acid medium with or without 10 mM glucose. A photo of a representative experiment is shown.

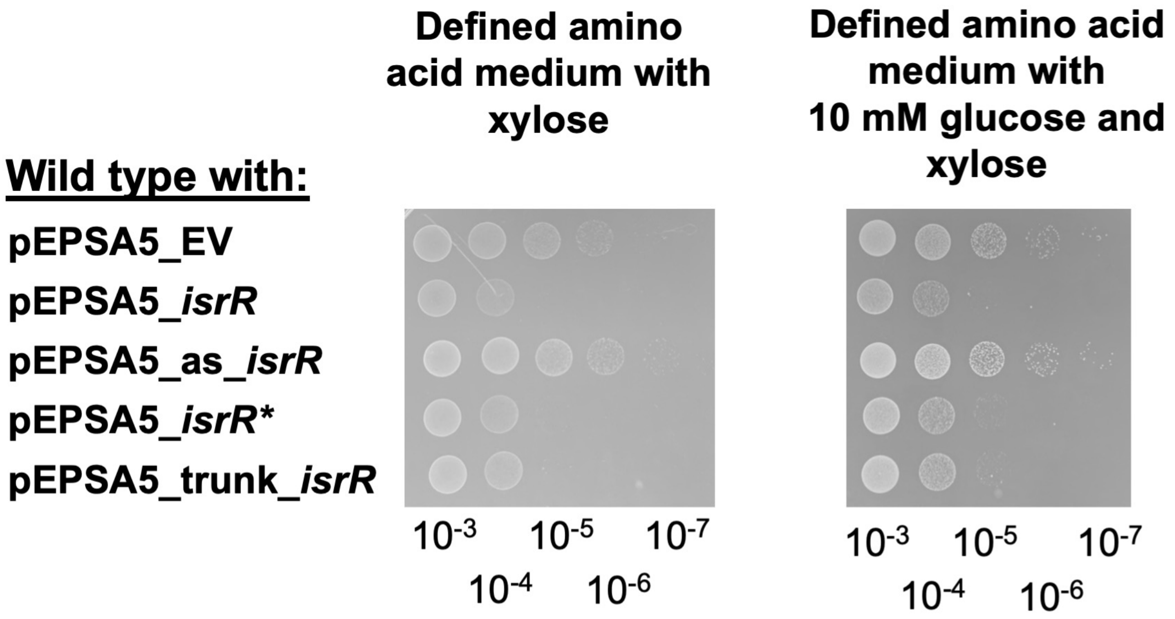


**Supplemental Figure 5.** Expression of *isrR* or *isrR** results in decreased growth on solid defined medium containing amino acids for carbon and energy. The wild type (JMB1100) containing pEPSA5, pEPSA5_*isrR,* pEPSA5_as_*isrR,* pEPSA5_*isrR**, or pEPSA5_trunk_*isrR* plasmids were standardized to an optical density ($A_{600}$) of 2, serial diluted (10-fold per dilution), and spotted on solid defined amino acid medium supplemented with chloramphenicol and 2% xylose with or without 10mM glucose. A photo of a representative experiment is shown.

**A**

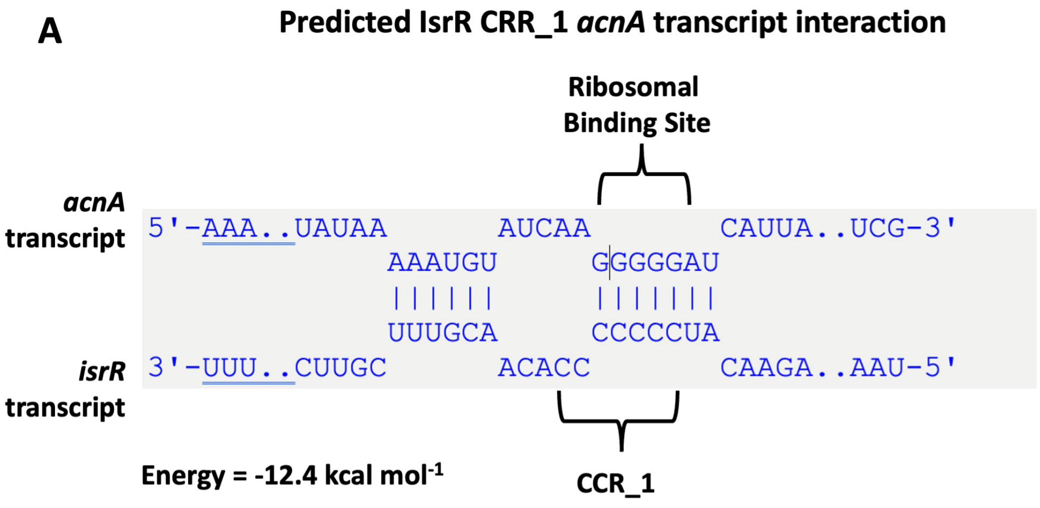


**B**

ATATAAAAATGTATCAAGGGGGATCATTAA**ATG**

Proposed interaction site highlighted.

**Supplemental Figure 6. Panel A.** IntaRNA predicted IsrR-*acnA* mRNA interaction. involving the first cytosine-rich region (CRR_1) of IsrR. **Panel B.** A portion of the 5' untranslated region (UTR) of the *acnA* transcript and the translational start codon of *acnA* ATG is shown (blue bold). The yellow highlight denotes predicted IsrR-*acnA* interacting nucleotides.

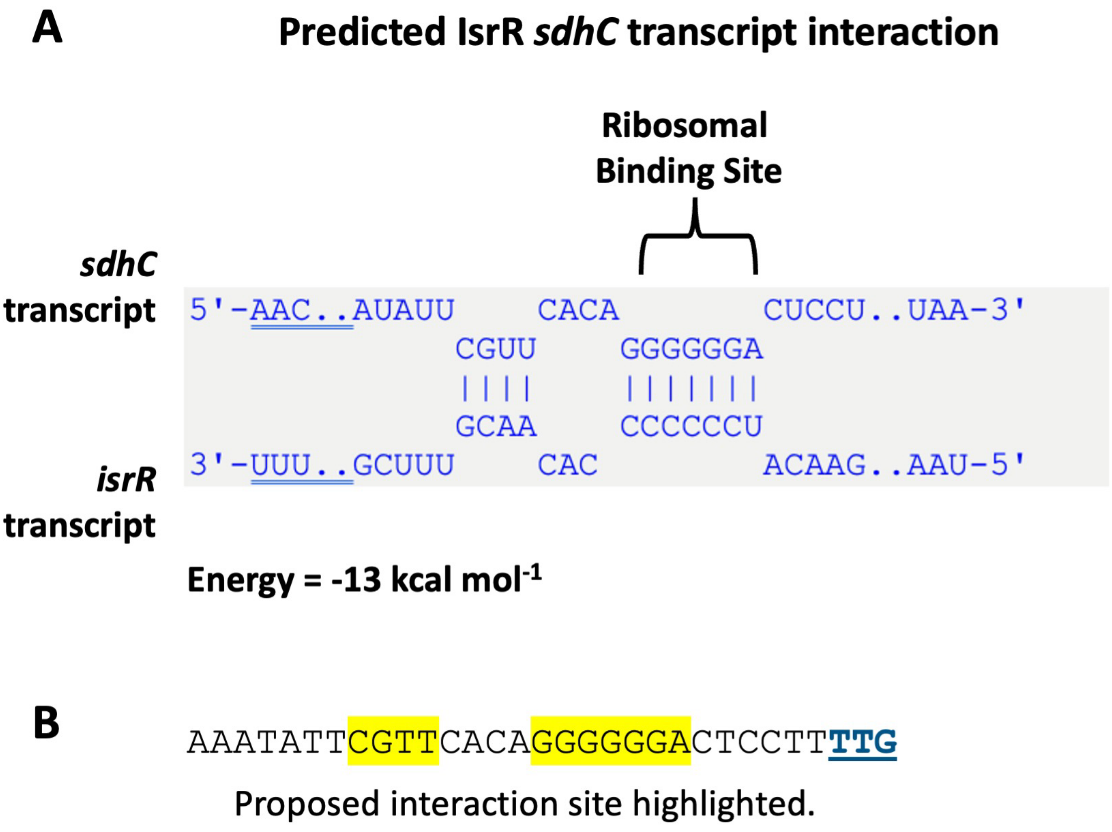


**Supplemental Figure 7. Panel A.** IntaRNA predicted IsrR-*sdhC* mRNA interaction. **Panel B.** A portion of the 5' untranslated region (UTR) of the *sdhC* transcript and the translational start codon of *sdhC* is shown (blue bold). The yellow highlight denotes predicted IsrR-*sdhC* interacting nucleotides.

**A**

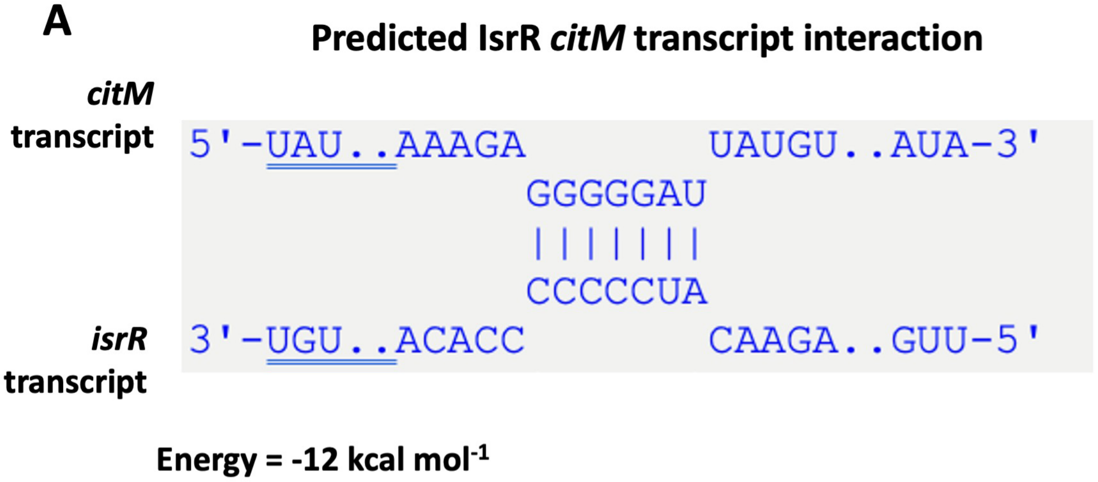


**B**

AGA==GGGGGAT==TATGTAAATTGTATTAAAAGTGGAGGGAGAAAATAAT**ATG**

Proposed interaction site highlighted.

**Supplemental Figure 8. Panel A.** IntaRNA predicted IsrR-*citM* mRNA interaction. **Panel B.** A portion of the 5' untranslated region (UTR) of the *citM* transcript and the translational start codon of *citM* is shown (blue bold). The yellow highlight denotes predicted IsrR-*citM* interacting nucleotides.

**A**

**Predicted IsrR *citZ* transcript interaction**

```
citZ
transcript   5'-UAA..ACAGU          AUUUA..UUU-3'
                      AAAGGGGAA
                      |||||||||
                      UUUCCCCUU
isrR         3'-UGU..UAUAU          AUAAA..GUU-5'
transcript
```

**Energy = -9 kcal mol$^{-1}$**

**B**

AAAGCCATTTCATAAAGAAACAGTAAAGGGGAAATTTATC**ATG**

Proposed interaction site highlighted.

**Supplemental Figure 9. Panel A.** IntaRNA predicted IsrR-*citZ* mRNA interaction. **Panel B.** A portion of the 5' untranslated region (UTR) of the *citZ* transcript and the translational start codon of *citZ* is shown (blue bold). The yellow highlight denotes predicted IsrR-*citZ* interacting nucleotides.

**A**

**Predicted IsrR *mqo* transcript interaction**

```
        mqo
 transcript 5'-UCU..UUUUA          CUG      UUA        CA     AUAGC..GAG-3'
                         AAAAGGGGGA   UAUUUG    UGAC AA   CAAC
                         ||||||||||   ||||||    |||| ||   ||||
                         UUUUCCCCUU   AUAAAC    ACUG UU   GUUG
       isrR 3'-UUU..GUAUA                   UUUU    G  AG     UUCUU..AAU-5'
 transcript
```

**Energy = -9.5 kcal mol$^{-1}$**

**B**

**ATG**TCGTATTTCTCGAACGTTCCATTATTCATTTTAAAAAGGGGGACTGTATTTGTTATGA
CAACACAACATAGCAAAACAG

Proposed interaction site highlighted.

**Supplemental Figure 10. Panel A.** IntaRNA predicted IsrR-*mqo* mRNA interaction. **Panel B.** A portion of the 5’ region of the *mqo* transcript and the translational start codon of *mqo* is shown (blue bold). The yellow highlight denotes predicted IsrR-*mqo* interacting nucleotides.

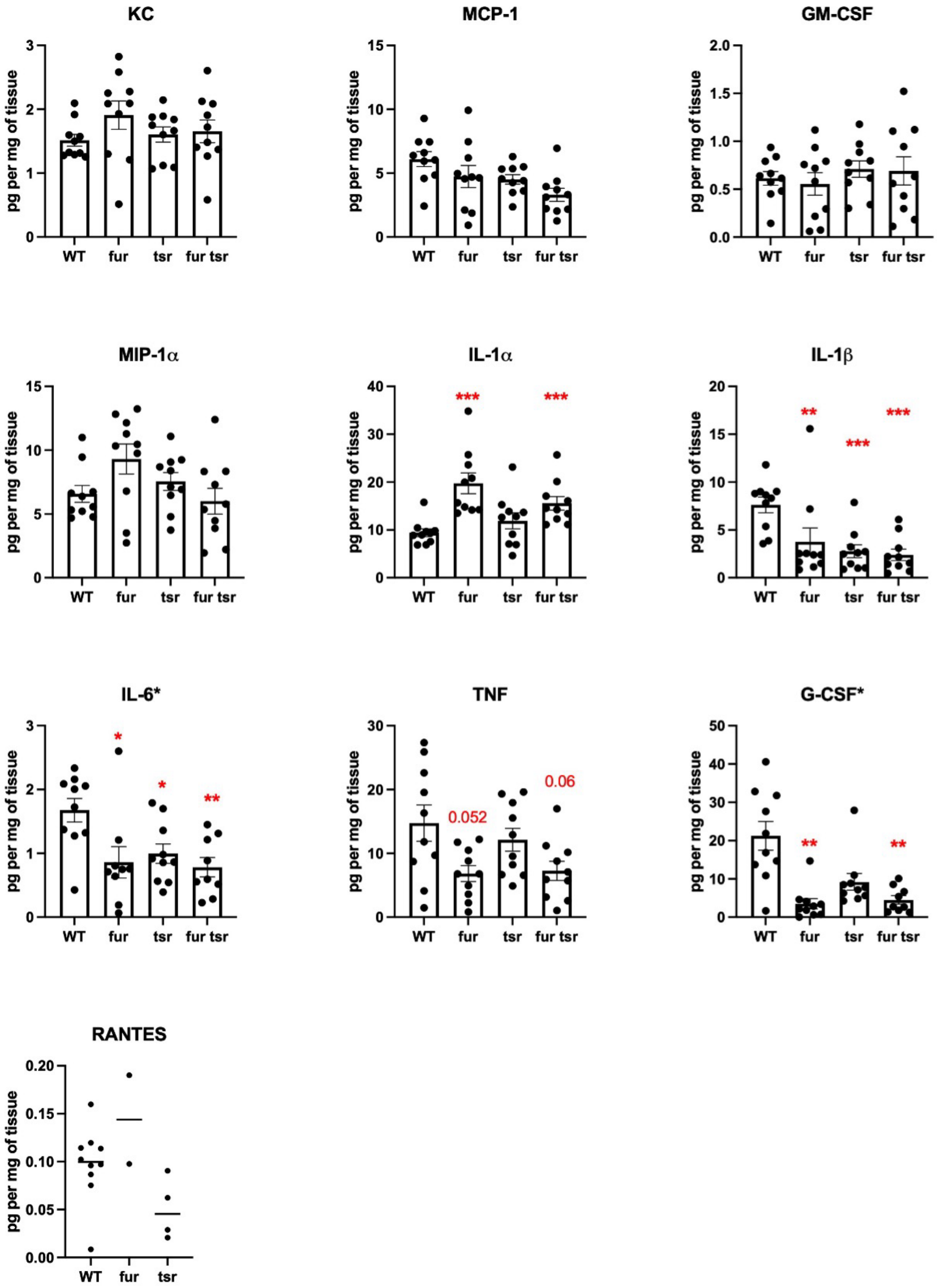


**Supplemental Figure 11.** Cytokine levels were quantified at four days post infection with wild type (WT) (JMB1100), Δ*fur::tetM* (JMB10842), Δ*isrR* (JMB11292), or Δ*fur::tetM* Δ*isrR* (JMB11293). Data is a representative experiment with n=10. Each dot is an individual animal, and the bar or line represents the mean. Error bars represent the SEM and may be smaller than symbols. * and ** indicates p<0.05 and p<0.01, respectively, compared to WT for Δ*fur::tetM* and Δ*fur::tetM* Δ*isrR* by Mann-Whitney test.
